# PLURIPOTENT STEM CELL-DERIVED CARDIOVASCULAR PROGENITORS DIFFFERENTIATED ON LAMININ 221 REGENERATE AND IMPROVE FUNCTION OF INFARCTED SWINE HEARTS

**DOI:** 10.1101/2021.04.29.441908

**Authors:** Lynn Yap, Li Yen Chong, Clarissa Tan, Swarnaseetha Adusumalli, Millie Seow, Jing Guo, Zuhua Cai, Sze Jie Loo, Eric Lim, Narayan Lath, Lei Ye, Enrico Petretto, Karl Tryggvason

## Abstract

**Background:** Ischemic heart disease is a huge global burden where patients often have irreversibly damaged heart muscle. State-of-the-art technology using stem cell-derived products for cellular therapy could potentially replace damaged heart muscle for regenerative cardiology.

**Methods and Results:** Pluripotent human embryonic stem cells (hESCs) were differentiated on a laminin LN521+221 matrix to cardiovascular progenitors (CVPs). Global transcriptome analyses at multiple time points by single-cell RNA-sequencing demonstrated high reproducibility (R^2^ > 0.95) between two hESCs lines. We identified several CVP signature genes as quality batch control parameters which are highly specific to our CVPs as compared to canonical cardiac progenitor genes. A total of 200 million CVPs were injected into the infarcted region caused by permanent ligation of the coronary arteries of 10 immunosuppressed pigs and maintained for 4- and 12-weeks post transplantation. The transplanted cells engrafted and proliferated in the infarcted area as indicated by IVIS imaging, histology staining and spatial transcriptomic analysis. Spatial transcriptomic analysis at 1 week following transplantation showed that the infarcted region expressed human genes in the same area as immunohistology sections. Heart function was analyzed by magnetic resonance imaging (MRI) and computerized tomography (CT). Functional studies revealed overall improvement in left ventricular ejection fraction by 21.35 ± 3.3 %, which was accompanied by significant improvements in ventricular wall thickness and wall motion, as well as a reduction in infarction size after CVP transplantation as compared to medium control pigs (P < 0.05). Immunohistology analysis revealed maturation of the CVPs to cardiomyocytes (CMs) where the human grafts aligned with host tissue forming end-to-end connections typical for heart muscle. Electrophysiology analyses revealed electric continuity between injected and host tissue CMs. Episodes of ventricular tachyarrhythmia (VT) over a period of 25 days developed in four pigs, one pig had persistent VT, while the rest remained in normal sinus rhythm. All ten pigs survived the experiment without any VT-related death.

**Conclusions:** We report a highly reproducible, chemically defined and fully humanized differentiation method of hESCs for the generation of potent CVPs. This method may pave the way for lasting stem cell therapy of myocardial infarction (MI) in humans.

**Clinical Perspective:** *What is New?:*

- We present a highly reproducible, chemically defined and fully humanized laminin-based differentiation method for generation of large amounts of cardiovascular progenitors (CVP); 20 million cells in a 10 cm^2^ culture dish which were used for a preclinical study in pigs.
- Transplantation of the CVPs into the myocardial infarcted pig hearts yields maturation of the progenitor cells to cardiomyocytes (CMs) and improved cardiac function (21.35 ± 3.3 % LVEF improvement) using only 200 million CVPs.
- Temporary episodes of ventricular arrhythmia (50%) were observed after CVP transplantation. No fatal ventricular arrhythmia occurred.

*What are the clinical implications?:*

- Our laminin-based approach generated potent CVPs *in vivo* and largely restored function of the damaged heart.
- Cardiovascular progenitors may provide a new and safe therapeutic strategy for myocardial infarction.
- The results may have a significant impact on regenerative cardiology.

## Introduction

Myocardial infarction (MI) is the most common cause of human death worldwide. Failure of damaged heart tissue to self-regenerate often leads to heart failure. Heart transplantation, although being curative, is constrained in its capabilities due to shortage of donor hearts. For over two decades, cell therapy approaches, including stem cell-based methods, have been considered as a possible solution for MI, but attempts to develop such treatments have thus far been unsuccessful. For example, clinical trials involving hypothetical endogenous heart tissue and mesenchymal stem cells have not yielded positive results^1–3^. Overall, inconsistency in stem cell differentiation methods and their variable reproducibility, together with large-scale falsifications of data have caused confusion and crisis in the field of regenerative cardiology^2^.

Delving deeper into studies concerning the use of stem cell-derived cells, in particular cardiomyocytes (CMs), and the transplantation of such cells is an alternative approach that holds promise for future regenerative cardiology. In a recent study, contractile CMs derived from pluripotent human embryonic stem cells (hESC) were transplanted into infarcted myocardia of pigs and nonhuman primates (NHP). While successful engraftment was observed in the infarcted host hearts, severe side effects in the form of graft-induced tachyarrhythmia and subsequent death predominantly hinder their use in human patients^4, 5^. Menasché’s group successfully placed a fibrin patch transplanted with CD15+ and ISL1+ progenitors into the epicardium of the infarcted area in humans^6, 7^, and arrhythmia was not detected. However, similarly to the two mentioned studies, the transplanted contractile CMs did not appear to improve heart function. Although the temporary nature of tachycardia might be managed with drugs or overdrive pacing, mature CMs have not proven effective in cardiac regeneration which underpins their unsuitability for use in cell therapy. Apart from CMs, it has also been shown that early ventricular progenitors expressing *ISL1,* differentiated from pluripotent stem cells, preserved heart function in infarcted murine heart^8^, suggesting applicability of cardiac progenitors over CMs in cell therapy of MI. However, as these progenitors were differentiated on mouse tumor extract Matrigel^®^, contamination with animal matrix deems these progenitors not optimal for clinical usage.

We have previously described a stable and highly reproducible method for differentiation of cardiovascular progenitors (CVPs) from pluripotent hESCs on a humanized matrix surface comprising the most abundant heart laminin, LN221^9^ that is present in the ultrathin basement membrane that surrounds single cardiac muscle fibers. These CVPs successfully engrafted and promote vasculogenesis in the infarcted murine heart tissue, ultimately forming sarcomeres as well as intercalated discs at end-to-end junctions^9^. Furthermore, the left ventricular ejection fraction was found to improve at 4- and 12-weeks following transplantation. These results, while promising, needed to be validated in a large animal model in detailed studies of cardiac muscle formation, heart rhythm and its functional effects. We have chosen pig as the model system since the pig heart is most physiologically similar to human heart^10^.

Our long-term goal is to explore the possibility of developing a viable cell therapy approach for human MI. In this study, we examined the therapeutic effects of 200 million non-contractile hESC-derived CVPs differentiated on the heart laminin LN221 matrix, and its regenerative potential on damaged heart tissue in a pig model after permanent ligation of the coronary arteries. In view of severe concerns surrounding transplantation of CMs, such as lethal arrhythmia and poor functional improvements^2^, we chose to transplant 11-day differentiated CVPs that previously was shown to generate well-structured cardiac fibers in mouse hearts^9^.

We propose that the present highly reproducible, chemically defined, humanized and ROCK inhibitor-free differentiation method can be used to produce clinical quality CVPs for treatment of MI. The CVPs generated in this manner are quality-controlled by detailed transcriptomics analyses, have shown to be relatively safe in pigs (all treated animals survived without tumor formation), and yield *in vivo* efficacy in large animal MI model. The CVPs described here will have a significant potential in the development of a cell therapy product for ischemic heart disease.

## Materials and Methods

The details concerning the materials and methods can be found in the online Data Supplements.

### Human Embryonic Stem Cells

All human pluripotent stem cell studies were carried out in accordance with National University of Singapore’s Institutional Review Board (IRB 12-451). Pluripotent hESCs H1 (karyotype: 46, XY; WiCell Research Institute, WA01) and HS1001 (karyotype: 46, XY, Karolinska Institute)^11^ were maintained on 10 μg/ml LN521 (Biolamina AB, LN521) coated culture plates with daily change of Nutristem^®^ (Biological Industries, 05-100-1A) medium. Routine monitoring of pluripotent markers (POU5F1 (Santa Cruz, sc-5279) and Tra1-60 (Millipore, MAB4360) by flow cytometry and genomic stability by karyotyping were performed at Singapore General Hospital cytogenetics laboratory.

On the day of differentiation (day 0), hESCs were seeded at 6 million cells into 10-cm^2^ dishes (ThermoFisher, 150464) coated with combination a matrix of (3.33 μg/ml) LN-521 and (10 μg/ml) LN-221 in 5ml of PBS. This protocol was based on our previous study^9^. At confluency (day 4), the cells were exposed to differentiation medium (RPMI 1640 (ThermoFisher, 11875-093) with B27 supplement without insulin (ThermoFisher, A1895601) and 10 μM of CHIR99021 (Tocris, 4423) for 24 hours. The next day (day 5), the medium was removed and replaced with differentiation medium. At day 7, the medium was changed to differentiation medium with the addition of 5 μM of IWP2 (Tocris, 3533) for 2 days. Wells were replaced with differentiation medium at day 9 and day 11. A 10-cm^2^ dish can generate ∼ 20 million cells. Therefore, to achieve 200 million cells for 1 pig, we performed differentiation in ten 10-cm^2^ dishes. Schematic presentation of the differentiation protocol is shown in figure 1A.

**Figure 1.**
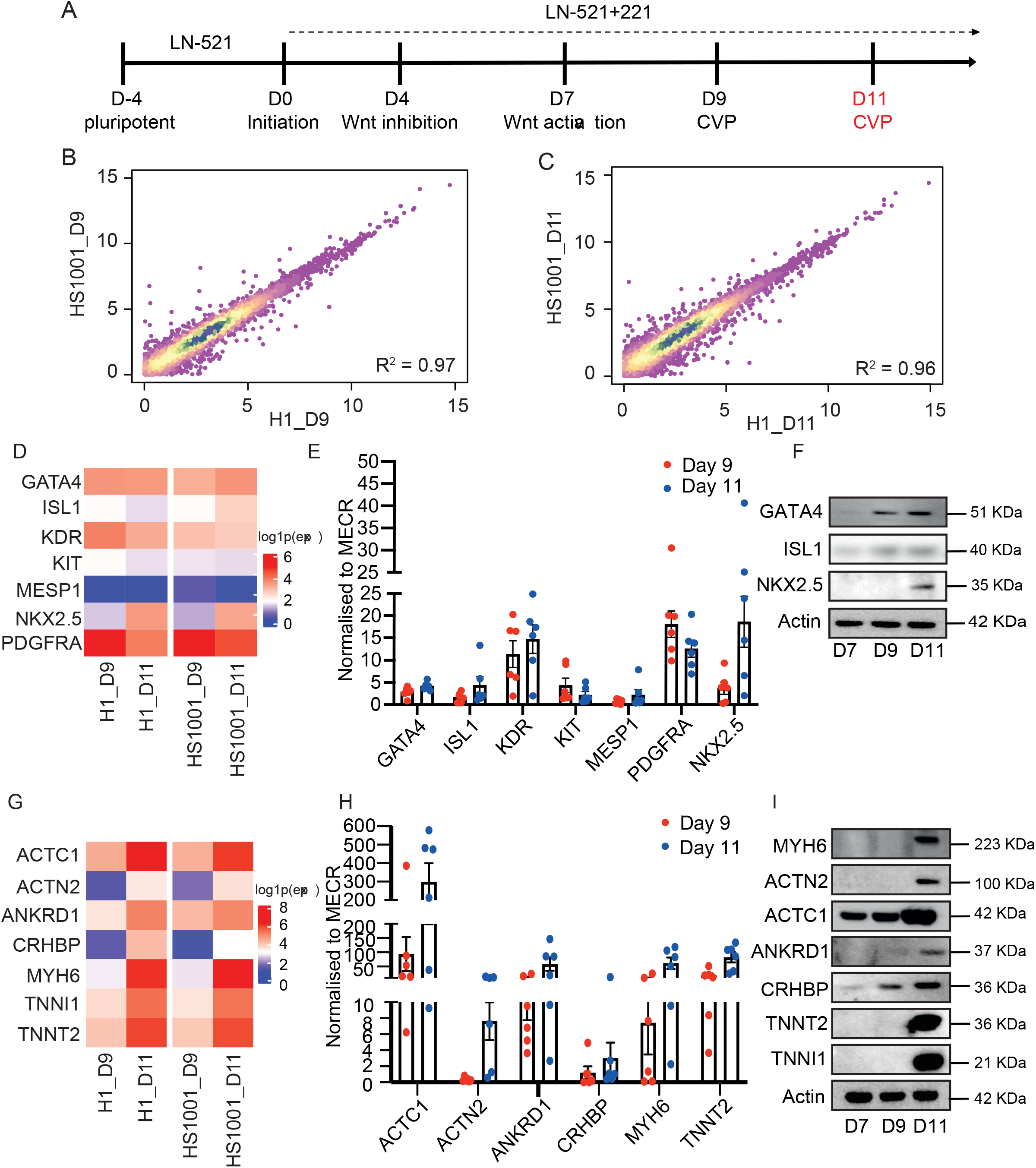
Differentiation reproducibility and hESC-derived cardiovascular progenitor (CVP) signature genes. **A)** Schematic timeline from pluripotent stem cells on LN521 matrix at day - 4 (D-4), day 0 (D0) initiation of differentiation on LN521+221 matrix, to day 4 (D4) Wnt signaling inhibition and day 7 (D7) Wnt signaling activation and finally towards day 9 (D9) and day 11 (D11) CVP formation. Day 11 CVP will be used in this study. **B-C)** Correlation plot of bulk RNA sequencing data from cell lines H1 and HS1001 hESCs at **(B)** day 9 and **(C)** day 11. There are high transcriptomic correlation between cell lines on day 9 (R^2^ = 0.97) and day 11 (R^2^ = 0.96). **D)** Heat map of canonical progenitor genes at differentiation days 9 and 11 in cell lines H1 and HS1001 (log1p transformed expression level). Supplementary table 1 shows the actual TPM for each gene. **E)** Quantitative PCR of canonical progenitor genes from differentiation days 9 and 11. The genes were normalized to MECR. There are no significant differences in all the groups (P > 0.05). Mean ± SEM, n = 6 individual differentiation batches. **F)** Immunoblotting of canonical progenitor genes from differentiation days 7, 9 and 11. Refer to supplementary figure 1 for the quantification of each blots. **G)** Heat map of the unique CVP signature genes at differentiation days 9 and 11 in cell lines H1 and HS1001 (log1p transformed expression level). Supplementary table 1 shows the actual TPM for each gene. **H)** Quantitative PCR results of day 11 CVP signature genes from differentiation days 9 and 11. The genes were normalized to MECR. Even though are no significant differences in all the groups (P > 0.05), there are a upward trend towards higher expression at day 11 as compared to day 9. Mean ± SEM, n = 6 individual differentiation batches. **I)** Immunoblotting of the day 11 CVP signature genes from differentiation days 7, 9 and 11. Refer to supplementary figure 1 for the quantification of each blots.

**Table 1.** Pigs characteristics and summary

| ID | Weight (kg) | Sex | Treatment | End-point | Electrophysiology analysis | Presence of VT? | CT findings |
| --- | --- | --- | --- | --- | --- | --- | --- |
| Ssc2667 | 10.3 | M | Sham | 12-week | 2 weeks |  | NA |
| Ssc2669 | 13.6 | F | Sham | 12-week | NA |  | NA |
| Ssc2676 | 13 | M | Sham | 12-week | NA |  | No tumors, simple cyst, one in either kidney |
| Ssc2515 | 13 | M | Media | 4-week | 4 weeks |  | NA |
| Ssc2545 | 12.6 | F | Media | 4-week | NA |  | NA |
| Ssc2548 | 12 | M | Media | 4-week | NA |  | NA |
| Ssc2666 | 10.6 | M | Media | 4-week | NA |  | NA |
| Ssc2668 | 11.8 | F | Media | 7-week | NA |  | NA |
| Ssc2665 | 10 | M | Media | 8-week | NA |  | NA |
| Ssc2585 | 11.8 | F | Media | 9-week | 2 weeks | No | NA |
| Ssc2517 | 15 | M | Media | 12-week | NA |  | NA |
| Ssc2572 | 11.6 | M | Media | 12-week | NA |  | 6 mm nodule in the right upper pulmonary lobe, no calcification, indetermined; 3.2x.2.8 mm expansile lytic lesion in mid sacrum, indetermined |
| Ssc2583 | 12.8 | M | Media | 12-week | 2 weeks | No | No tumors |
| Ssc2469 | 10.82 | F | D11 | 4-week | NA | No | NA |
| Ssc2520 | 13 | M | D11 | 4-week | NA | No | NA |
| Ssc2584 | 13.1 | F | D11 | 4-week | 2 weeks | No | NA |
| Ssc2578 | 11.86 | M | D11 | 10-week | NA | Yes | NA |
| Ssc2576 | 12 | M | D11 | 10-week | NA | Yes | No tumors |
| Ssc2498 | 13.4 | F | D11 | 12-week | NA | Yes | No tumors |
| Ssc2519 | 13.6 | M | D11 | 12-week | 4 weeks | No | No tumors |
| Ssc2577 | 12.8 | M | D11 | 12-week | 2 weeks | Yes | No tumors |
| Ssc2573 | 12 | F | D11 | 12-week | 2 weeks | Yes | No tumors |
| Ssc2673 | 12.6 | M | D11 | 12-week | NA | No | No tumors |

### Myocardial Infarction Model in Pigs

Either gender of 3-month-old Sus scrofa pigs at 13-15 kg were purchased from SingHealth Experimental Medicine Centre (SEMC, Singapore) and used in all experiments. Immunosuppression was given to prevent rejection of the human cells in the pig’s heart. Five days before the surgery, cyclosporine (Novartis, ADP835296) was given in the diet twice daily at 15 mg/kg and maintained throughout the experiment. First dose of abatacept (Orencia^®^, Bristol-Myers Squibb, ABT4318) at 12.5 mg/kg was given on the day of surgery via intravenous injection and once every 2 weeks until the end of the experiment. Corticosteroid immunosuppression using methylprednisolone (*Vem ìlaç*, 011001) was administered intraperitoneally once daily (250mg on surgery day, 125 mg for first 2 weeks after transplant and 62.5 mg until the end of the experiment). All the procedures of animal handling were performed with prior approval and in accordance with the protocols and guidelines of SingHealth’s Institutional Animal Care and Use Committee (IACUC) (2018/SHS/1426).

On the day of surgery, the pigs were randomly assigned to 3 groups: sham pigs are healthy controls without MI (12-weeks, n = 3), MI pigs injected with medium only (4-weeks, n = 4, 12-weeks, n = 6), MI pigs transplanted with day 11 CVPs (4-weeks, n = 3, 12-weeks, n = 7). MI was generated by permanent ligation of the left anterior descending coronary (LAD) and left circumflex (LCX) arteries, followed by intramyocardial injection of 200 million luciferase-labeled CVPs into the infarcted region.

Magnetic resonance imaging (MRI) was performed at 1, 4 and 12-weeks post-transplantation to determine the cardiac function. Computerized tomography (CT) scans were carried out at 12-weeks to identify potential tumor development in the CVP-transplanted animals as a safety study. Electromechanical mappings using the 3D-NOGA system were performed at weeks 2 and 4 for assessment of myocardial viability and to localize possible arrhythmogenic foci. Implantable cardiac monitors (Medtronic plc, Minneapolis, MN, USA) were inserted for continuous single-channel electrocardiogram (ECG) monitoring. At the end of the experiment, the pig hearts were imaged with an IVIS Spectrum imaging platform (Perkin Elmer) to determine the location of the human grafts and subsequent downstream tissue processing and histology. Schematic presentation of the immunosuppression and surgery regimen is shown in figure 3A.

## Results

### Differentiation Reproducibility and hESC-derived Cardiovascular Progenitor (CVP) Signature Genes

Since therapeutic hESC-derived cells are defined as drugs by the FDA and EMA^12^, we set out to explore if the CVPs generated and used here share homogeneous phenotypic characteristics, and fulfil acceptable safety and efficacy criteria, i.e. similar transcriptome biomarkers, absence of tumorigenicity, in vivo regeneration and clinical effects as we have previously reported for mice^9^.

We first characterized the transcriptome biomarkers of our CVPs. Pluripotent stem cells derived from two separate hESC lines, H1 and HS1001, were maintained in culture on a LN521 coating. These cells show strong expression of pluripotency markers Oct3/4 and Tra1-60 **(Supplementary Figure 1A)** and were transferred on day 0 to culture plates coated with a mixture of LN521 and LN221 for expansion and differentiation to CVPs as previously reported^9^ and as depicted in Figure 1A. Analysis of the expressed genes at days 9 and 11 showed high correlation of the expression profiles of two hESC-lines (day 9: R^2^ > 0.97, day 11: R^2^ > 0.96) (Figure 1B and C). This demonstrates the high transcriptional reproducibility and stability of the differentiated LN221-dependent stem cells, which underpins their applicability as potential therapeutic drugs.

To further understand and characterize the cells with the goal of defining a set of CVP signature genes (at day 11) for quality control, we performed the differential gene expression between differentiation days 9 and 11 to obtain a set of expressed genes typical for day 11 CVPs. First, we examined canonical, previously reported progenitor genes (*GATA4*^13^, *ISL1*^14^, *KDR*^15^, *KIT*^16^, *MESP1*^5^, *NKX2.5*^17^ and *PDGFRA*^18^) in our days 9 and 11 CVPs (Figure 1D). Expression of these genes was highly similar between the two cell lines, again supporting the reproducibility and stability of the differentiated cells observed by global transcriptomes^9^ (Figure 1D). However, we observed that the day 11 CVPs had low expression of the canonical progenitor genes. For example, genes *ISL1*, *KDR*, *KIT, MESP1* and *PDGFRA* have low expression at day 11 **(Supplementary Table 1)**. Other genes *GATA4*, *NKX2.5*, showed higher expression at day 11 as compared to day 9 which could be used to identify our cells. We then went about to validate the RNAseq results with quantitative PCR (qPCR) (Figure 1E), and also quantified the protein levels by immunoblotting (Figure 1F **and Supplementary 1B)**. Both qPCR and immunoblotting results confirmed that these known genes had variable low expressions and, thus, are not ideal markers for a CVP population (except for NKX2.5 and GATA4) to be used in reconstruction of the damaged heart muscle. Therefore, we went on to search for an additional set of more biologically meaningful gene markers that are better suitable for characterization of these CVPs.

To identify unique CVP signature genes may better describe a cell population with biological usefulness, we analyzed time-course bulk RNAseq and identified CVP markers that clearly distinguish day 9 vs day 11 compared to other timepoints. These signature genes include *ACTC1, ACTN2, ANKRD1, CRHBP, MYH6, TNNT2* and *TNNI1*. These genes were highly expressed and suggested biological relevance to cardiac muscle development at day 11 as compared to day 9 in both H1 and HS1001 (Figure 1G**, Supplementary Table 1)**. We validated the RNAseq results by comparing the expression levels of these signature genes from different differentiation batches using qPCR (Figure 1H), immunoblotting (Figure 1I) and flow cytometry **(Supplementary Figure 1D),** all showed gradual increase in expression at day 11 compared to day 9. Densitometry measurements of the immunoblot in figure 1I is shown in **Supplementary figure 1C**. Even though the statistical analyses between these 2 days from the qPCR data (Figure 1H) did not reveal significant differences (P> 0.05), we observed a consistent upward trend of gene expression on day 11. Expression of these biomarkers can be considered representative for genes at day 11 CVPs and we used it in every differentiation batch as quality control criteria. These cells were then used for the subsequent *in vivo* transplantation following MI.

### Spatial Transcriptomic Revealed Engraftment of Human Cells

To confirm the engraftment of the CVPs into the infarcted heart of the pigs, we performed spatial transcriptomics. The LAD and LCX were permanently ligated and 200 million CVPs (n = 2, replicate-1 and replicate-2) or medium (n = 1) were transplanted into the infarcted region. A healthy pig heart was used as a sham control (n = 1). One week after surgery, the animals were euthanized and the left ventricle was sectioned into 5 cross-sectioned rings (Figure 2A). The rings were imaged under the IVIS machine and tissues with positive luciferase signal were sectioned into smaller pieces. The pieces were analyzed with 10x Visium Spatial Gene Expression kit.

**Figure 2.**
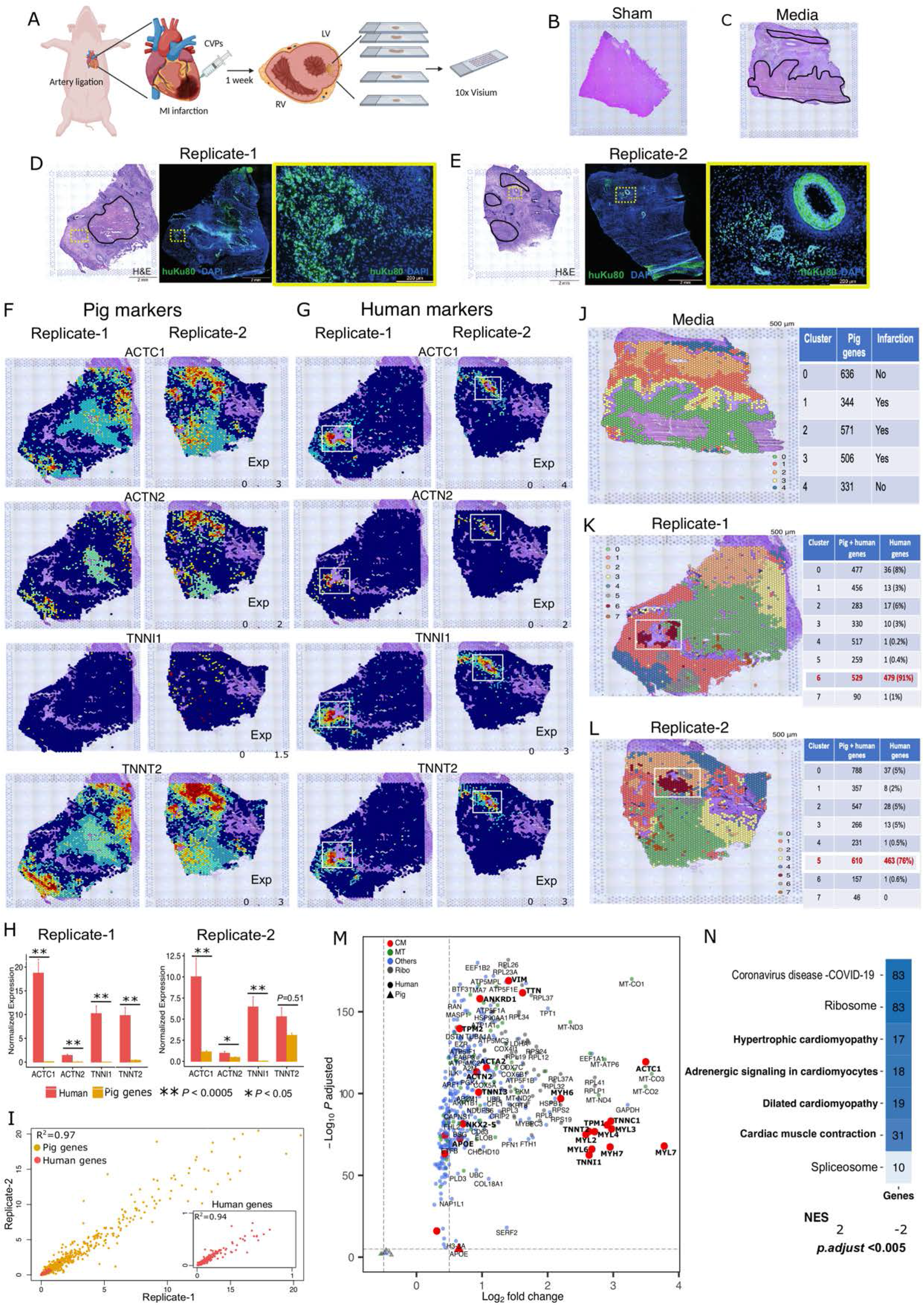
Spatial transcriptomic revealed engraftment of human cells. **A)** Schematic of four individual pig heart ventricular sections including a pair of ‘”biological” replicates’ with human CVPs engrafted. H&E staining and tissue section imaging was completed as described in the 10x Visium protocol. **B-C)** H&E staining images of **(B)** sham and **(C)** medium control tissue sections. Healthy tissues are demarcated with a black line in **(C)** and the rest of the tissues are infarcted regions. **D-E)** Histology staining of **(D)** replicate-1 and **(E)** replicate-2 (Left) H&E images of replicates with human CVPs engrafted at the highlighted regions in yellow. Healthy tissues are demarcated with a black line and the rest of the tissues are infarcted regions. (Middle) immunofluorescence images of engrafted human cells identified using human specific Ku80 (huKu80) antibody (green). (Right) Insert shows the higher magnification of engrafted human cells in the yellow box from the middle image. **F-G)** Spotplots with H&E images and spot overlays depicts the log-transformed normalized expression for pig and human *ACTC1, ACTN2, TNNI1* and *TNNT2* marker genes. The white box shows regions with human CVPs engraftment. **H)** Barplots shows quantified expressions of pig and human gene in the human cells engrafted spots (white box highlighted in panel F-G). T-test, *P*-value < 0.05 *; *P*-value < 0.0005***. Error bars represent standard error of the mean (SEM). **I)** Correlation plot of all genes including pig and human genes from biological replicates with engrafted human cells (R^2^ = 0.97). Insert shows the correlation of human genes in engrafted regions (white box) of replicates (R = 0.94). **J-L)** Spotplots with H&E images and spot overlays with unbiased clustering of spots based on global gene expression within individual spots in **(J)** medium, **(K)** replicate-1 and **(L)** replicate-2. Scale bar = 500 μm. The white box shows regions with human CVPs engraftment. The tables on the right shows the number of significantly expressed pig and human genes in each cluster post quality control (adjusted *P*-values < 0.05). **M)** Volcano plot of common human genes expressed in clusters specific to engrafted regions (white box) from the replicates (clusters 6 and from biological replicates 1 and 2 respectively). All genes are significantly differentially expressed with adjusted *P*-values < 0.05. **N)** Functional processes enriched in common human genes from biological replicates were computed by gene set enrichment analysis (GSEA) (adjusted *P*-values < 0.05). Normalized enrichment scores (NES) denote the upregulation and downregulation enrichment strength. The numbers in heatmap indicates the number of genes enriched for each function.

H&E staining was performed on the tissues to determine their tissue condition (Figure 2B - E). Based on intensity of the stains, we determined that all sham tissues were healthy (Figure 2B), medium control had regions of infarction and non-infarction (black outline) (Figure 2C) and replicates-1 and −2 had regions of infarction, non-infarction (black outline) and human cells engrafted (outlined in yellow box) (Figure 2D and E, left). Next, we stained the tissue with human specific anti-Ku80 to confirm the position of the engrafted regions and identified the presence of human cells (Ku80+, green) (Figure 2D and E, middle, yellow box). A magnified view of the engrafted region (yellow box) is shown in figure 2D and E, right panel.

The spatial transcriptomic (ST) reads (from healthy, medium control, CVP replicate-1 and CVP replicate-2 sections) were aligned to the pig and human combined genome references to segregate the reads belonging to the transplanted CVPs from the host pig heart reads. After mapping and quality control, we obtained gene-spot matrix consisting of 11328 genes and 9269 individual capture locations (spots) on the ST array (summarized over four sections). We then subjected the data to downstream computational analysis. The sham, medium, CVPs replicate-1 and −2 sections covered 2187, 2087, 2948 and 2047 spots under the tissue. To confirm the successful engraftment of the CVPs in the replicate-1 and −2, we assessed the expression of markers *ACTC1, ACTN2 TNNI1, TNNT2* (reported in Figure 1G) and other CVP markers, such as *MYH7, MYL2, TNNC1 and TPM1* in the engrafted regions (Figure 2F - G**, Supplementary figure 2A - B**). We showed the human markers were indeed specifically expressed in the engrafted region as highlighted by H&E staining, suggesting successful CVPs engraftment (Figure 2G). Subsequently, we quantified the expression levels of the pig and human markers in the engrafted regions in each replicate (Figure 2H**, Supplementary figure 2C**). We observed significantly higher expression of human CVP markers *ACTC1, ACTN2, TNNI1* compared to those from pigs in both replicates (*P* < 0.05), whereas human *TNNT2* is significantly highly expressed in replicate-1 (Figure 2H). We then sought to assess the transcriptional similarity across the CVP-engrafted replicates and performed correlation analysis across all genes from tissue sections and human genes in the engrafted regions. The correlation analysis revealed high transcriptional reproducibility between replicates (Figure 2I, Spearman’s correlation level of 0.97 and 0.94 for all genes and human genes, respectively).

Furthermore, to identify subgroups of spots with similar gene expression profiles and spots underlying the human CVP-engrafted regions by data driven approach, we performed unsupervised clustering. We embedded the data into low-dimensional space via non-negative matrix factorization and performed clustering independently on each section using a graph-based approach. To characterize the clusters into context and assess their spatial organization, spots were projected on the H&E-stained tissue images. We identified 6 clusters in sham, 5 clusters in medium and 8 clusters each for replicate-1 and −2 (Figure 2J - L**, Supplementary figure 2D)**. The differentially expressed genes across 5 sham clusters were enriched for different signaling pathways and diabetic cardiomyopathy (**Supplementary figure 2E**). Clusters 0 and 4 from the medium sections were specific to the non-infarcted regions (Figure 2J) whereas, the other clusters were associated with the spots in the infarcted regions. To characterize molecular functions and pathways within the non-infarcted and the infarcted regions, the differential gene expression analysis was performed. We observed that the non-infarcted regions were highly enriched for genes associated with cardiac function including cardiac muscle contraction, dilated cardiomyopathy, adrenergic signaling in CMs and others (adjusted *P* < 0.05). In contrast, genes upregulated in infarcted regions were enriched for pathways such as glycolysis/gluconeogenesis, ECM-receptor interaction, human papillomavirus infection (**Supplementary figure 2F**).

We further sought to characterize spots with engrafted CVPs in replicate-1. The clustering analysis revealed that the majority of cluster 6 expresses the human genes, highlighting the CVPs engrafted region (Figure 2K). The differential gene expression analysis identified 529 genes that were significantly expressed in cluster 6 compared to all other clusters. About 91 % (479/529) of the differentially expressed (DE) genes in cluster 6 were human genes that are all upregulated (Figure 2K**, right**). Of the 479 human genes expressed, 5 % (26 genes) were cardiac markers that include *ANKRD1*, actinin-, myosin-chain genes, titin, troponin and vimentin genes. Interestingly, we observed about 29 % of human genes expressed to be mitochondrial (n = 55) and ribosomal (n = 80) genes as hESCs undergo mitochondrial biogenesis and activation of oxidative phosphorylation as key regulatory events in cell differentiation to beating CMs^19^. The remaining 66 % human genes include various genes such as collagens, laminins, heterogeneous nuclear ribonucleoproteins and others. The remaining 9 % (50/529) of the DE genes from cluster 6 are pig marker genes, of which 96 % (48/50) were downregulated, including *MYH7, S100A11* and *TIMP1*. Consistent with replicate-1, the clustering analysis in replicate-2 demonstrated a unique cluster (cluster 5) expressing majority of human genes (Figure 2L). About 76 % (463/610) of cluster 5 DE genes were from engrafted cells (Figure 2L**, right**). Similarly to replicate-1, the cardiac, mitochondrial and ribosomal genes constitute to 5 % (24 genes) and 28 % (51 and 80 mitochondrial and ribosomal genes) respectively, of the human genes from replicate-2. In contrast, engrafted cluster 5 from replicate-2 constitutes about 24% of pig DE genes (147/610), including cardiac markers such as *ACTA2, ACTG1, MYL2, VIM* and *TPM4* which were downregulated (log2 fold change < −0.5).

We next examined the common human markers from engrafted clusters from both replicates and found that about 97% of the marker genes were reproduced in both replicates (Figure 2M). This result demonstrates the reproducibility of engraftment between replicates consistent with the correlation analysis (Figure 2I inner panel). The functional analysis on the common human genes from engrafted clusters revealed significant enrichment of cardiac pathways including hypertrophic cardiomyopathy, adrenergic signaling in cardiomyocytes, dilated cardiomyopathy and cardiac muscle functions (adjusted *P* < 0.05) (Figure 2N).

Taken together, the ST results suggest successful engraftment of human CVPs in the infarcted host pig hearts. We showed significant expression of key human CVP marker genes, such as *ACTC1, ACTN2, TNNI1*, and *TNNT2* and others 1-week post transplantation. We also demonstrated high reproducibility of engraftment across biological replicates.

### Human CVPs Engraft and Form Mature CMs in Infarcted Pig Heart and Display Normal Muscle Histology in Highly Vascularized Human Grafts

All the pigs received immunosuppression five days before surgery and the immunosuppression was maintained throughout the 4- or 12-week long experiments. Magnetic resonance imaging (MRI) was performed at 1, 4 and 12 weeks after transplantation to assess heart function, and computerized tomography (CT) scan was done at 12-weeks to identify any potential tumor development in the pigs as a long-term safety study (Figure 3A **and Supplementary table 3)**. On the day of surgery, we generated infarction in the left ventricle by permanently ligating the arteries in an open chest surgery. Following the MI, we injected a total of 200 million CVPs in 1 ml of medium to approximately 10 sites intramyocardially into the macroscopically visible peri-infarct and infarcted regions. The animals were subsequently maintained for 4- and 12-week periods for observation **(Table 1)**. We confirmed the immunosuppression to be adequate based on immunohistology staining of the human muscle grafts showing the absence of immune cells with pan-leukocytes (CD45), B-lymphocytes (CD20) and T-lymphocytes (CD3) **(Supplementary figure 2)**. However, 3 pigs from the medium control group and 2 pigs from the CVP transplanted group did not manage to reach the endpoints due to complications from immunosuppression (anemia or lung infection). Refer to **table 1** and **supplementary table 3**.

**Figure 3.**
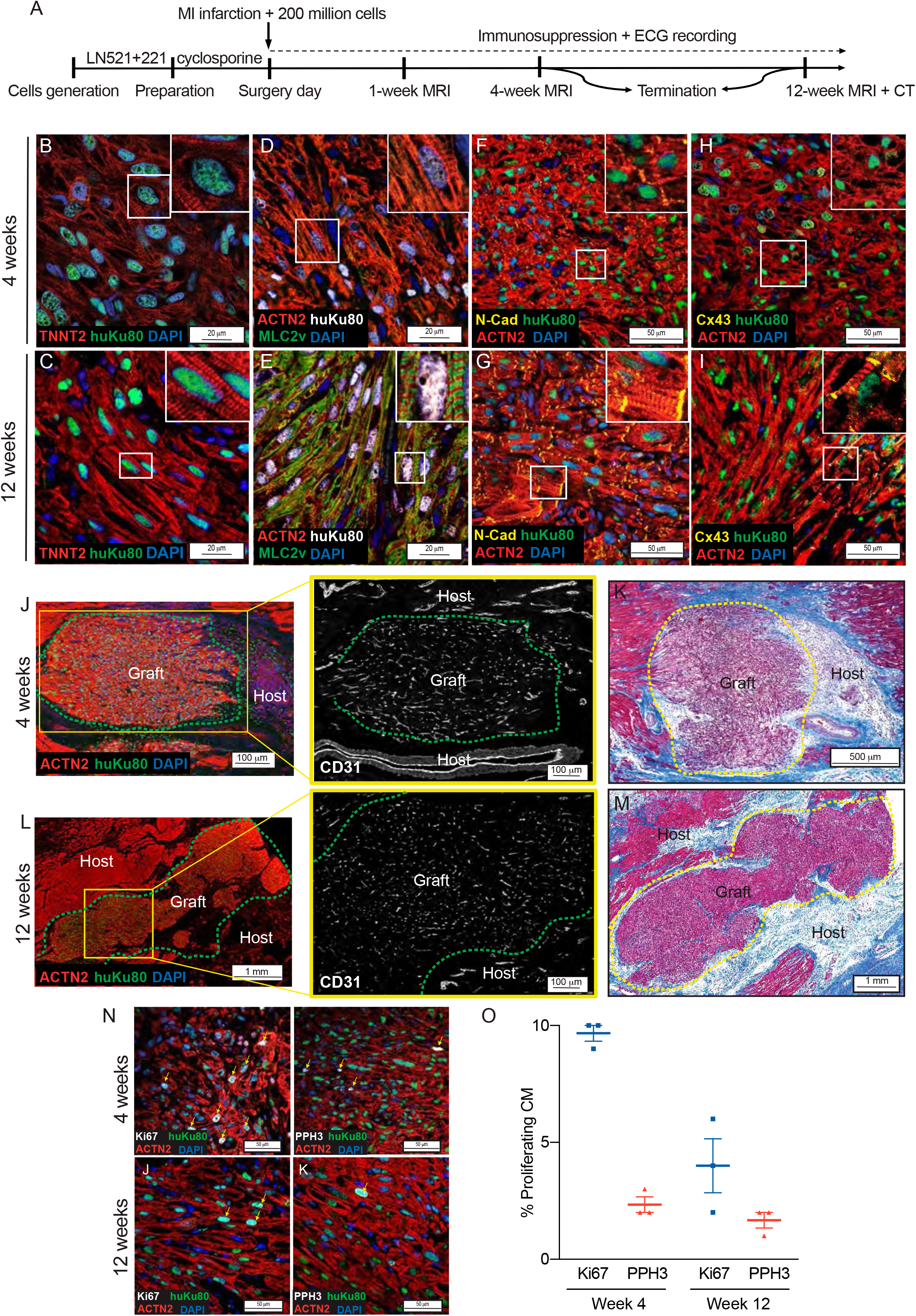
Human CVPs engraft and form mature CMs in infarcted pig heart and display normal muscle histology in highly vascularized human grafts. The engrafted CVPs matured in the scar and initiated growth of human heart muscle well vascularized by blood vessels. **A)** Schematic to depicts the timeline of events in pigs from CVP generation to permanent MI infarction in pigs and intramyocardial transplantation of CVP to post-transplantation MRI and CT scans and finally euthanization at 4- or 12-weeks post transplantation. The pigs were immunosuppressed and implanted with a cardiac monitor to continuously record the ECG throughout the experiment. **B, D, F and H)** Confocal immunofluorescence staining of 4-weeks post-transplanted grafts with specific antibodies. Insert shows higher magnification. Human cells were identified using human specific Ku80 (huKu80) antibody. At 4-weeks, the human grafts showed disorganized staining of **(B)** TNNT2 (red), **(D)** ACTN2 (red) and MLC2v (green), **(F)** N-Cadherin (N-cad, yellow) and ACTN2 (red) and **(H)** connexin-43 (Cx43, yellow) and ACTN2 (red). All nuclei are stained with DAPI (blue). **C, E, G and I)** Confocal immunofluorescence staining of 12-weeks post-transplantation grafts with specific antibodies. Insert shows the higher magnification. Human cells were identified using human specific Ku80 (huKu80) antibody. Human grafts shows highly organized staining of **(C)** TNNT2 (red), **(E)** ACTN2 (red) and MLC2v (green), **(G)** N-Cad (yellow) and ACTN2 (red) and **(I)** Cx43 (yellow) and ACTN2 (red). Highly organized mature striation were imaged at 12-weeks. All nuclei are stained with DAPI (blue). **J)** Left: Confocal immunofluorescence staining of 4 weeks post-transplantation grafts with ACTN2 (red), huKu80 (green) and DAPI (blue). The length of the graft was approximately 1 mm. Right: A zoom in image of the yellow box showing anti-CD31 (white) staining. The human graft is highly vascularized with many small blood vessels in and around the grafts. Green outline marks the human graft and host tissue surrounding the graft. **K)** Masson trichrome staining of the transplanted region in **(J)** shows low amount of collagen deposition (blue) around the human graft (purple) and host tissue at the edges (dark red). Yellow outline demarcated the human graft. **L)** Left: Confocal immunofluorescence staining of 12-weeks post-transplantation grafts with ACTN2 (red), huKu80 (green) and DAPI (blue). The length of the whole graft was approximately 5 mm, a significant increase in length as compared to 4-weeks. Right: A zoom in image in the yellow box showing anti-CD31 (white) staining. The human graft are similarly vascularized with small blood vessels in and around the transplantation area. Green outline marks the human graft and host tissue surrounding the graft. **M)** Masson trichrome staining of the transplanted region showing low amount of collagen deposition (blue) around the human graft (purple) and host tissue at the edges (dark red). Yellow outline marks the human graft. Refer to supplementary figure 4 for other H&E images of 12-weeks human grafts. **N)** Confocal immunofluorescence staining of 4- and 12-week human grafts with proliferation markers Ki67 or PPH3 (yellow arrows). **O)** Quantification of images in **(N)** reflecting the percentage of proliferating human CM expressing Ki67 (9.7 ± 0.3 %) and PPH3 (4 ± 1.2 %). There was a reduction in proliferative potential of the cells overtime. Mean ± SEM, n = 3.

At the end of 12 weeks, the pigs were euthanized, and the hearts were excised. The ligation sutures were visible and intact from the day of euthanization **(Supplementary figure 3A)**. Viability of the transplanted cells in the infarction site could also be monitored using live *in vivo* IVIS imaging following administration of luciferin into the coronary artery at the time of euthanization **(Supplementary figure 3B)**. Areas of heart muscle fibers with positive luciferase signal were identified and processed for histology staining. Hematoxylin and Eosin (H&E) staining showed large human grafts in close proximity to the normal pig muscle tissue in the infarcted region **(Supplementary figure 3C)**. Identity of human grafts was confirmed by immunohistology staining with a human nucleus specific antibody (anti-Ku80), cardiac troponin T (TNNT2), cardiac troponin I (TNNI3) alpha-actinin (ACTN2) and myosin light chain 2V (MLC2V) at week 4 and 12 (Figures 3B to E **and Supplementary figure 4A - B)**. We observed Ku80 positive (Ku80+) human cardiac muscle grafts with highly organized sarcomere striations as the graft mature from 4 to 12 weeks. We observed alternate ACTN2 and MLC2V striations at 12 weeks (Figure 3E) but these were initially absent at 4 weeks (Figure 3D). It is noteworthy that almost all the human cells expressed ventricular-like MLC2V instead of the atrial-like MLC2a **(Supplementary figure 4C - D)**, suggesting that most of the cells are ventricular but not atrial CMs.

Following identification of the human graft, we continued to investigate the presence of cell-to-cell connections with N-Cadherin (N-Cad) which anchors the myofibrils with connexin-43 (Cx43), a major electrical determinant of the electrical properties of the gap junctions of human CMs (Figure 3F - I). We observed only weak expression and spotty distribution of N-Cad staining at 4 weeks, but the proteins became better organized and highly expressed at 12-weeks. Moreover, we observed similar but weaker Cx43 expression at 4 weeks. Thereafter, the Cx43 exhibited stronger expression and highly organized distribution by 12 weeks. As Cx43 regulates intercellular coupling and conduction velocity, these results suggested that the human grafts continue to mature and become electrically coupled and physiologically functional over time.

We observed rapid engraftment, with extensive new heart tissue growth and functional development as early as at 4 weeks. This regeneration process cannot occur without sufficient blood supply. We further investigated whether the human grafts were vascularized by staining the tissue with CD31 (Figure 3J and K). The human graft stained positive for CD31 as early as at 4 weeks post-transplantation. Since these blood vessels were not positive for human-Ku80, it is likely that they were derived from the collateral blood vessel network of the myocardium surrounding the scar. We observed scarring in the infarcted region with Masson Trichrome staining. The results showed that the area surrounding the scar has lesser collagen deposition (Figure 3K and M) as compared to medium control hearts **(Supplementary figure 5)**. The extensive growth of micro-vessels even when the LAD and LCX were totally ligated and with reduction in fibrosis implies that these CVPs have chemoattractant properties that encourage the growth of new blood vessels and reduced scarring.

Since our CVPs are highly proliferative, we performed immunohistology staining for proliferation markers, Ki67 and phophohistone H3 (PPH3) to quantify the proportion of proliferating cells at 4 and 12-weeks after transplantation (Figure 3N and O). At 4 weeks, 9.7 % ± 0.3 % of the cells are positive for Ki67 and about 2.3 % ± 0.3 % were positive for PPH3 (P<0.05). In contrast, there was a significant reduction in the CVP proliferation rate of only 4 % ± 1.2 % being Ki67 positive and 1.7 % ± 0.3 % were positive for PPH3 at 12 weeks. The results indicate that the cells are highly proliferative at 4 weeks, but that they are already exiting the cell cycle at 12 weeks and becoming terminally differentiated CMs in the heart muscle where mature well-connected human cardiac fibers were formed (Figure 3L).

### Regenerated Infarcted Pig Heart Yields Concomitant Improvement in Heart Function

At the end of the 12-week timeline, the pigs underwent MRI and CT scans to determine heart function and long-term safety profile (n = 9-10 at weeks 1 and 4, n = 3-5 at week 12) **(Supplementary table 3)**. The left ventricles (LV) were cross-sectioned into 5 rings from the apex to the atrium. We observed gross morphology differences in the sham, medium and CVP transplanted hearts (Figure 4A). Sham had uninjured LV with healthy ventricular walls, medium control showed transmural scar and adverse ventricular remodeling (red arrows) in the lateral LV wall, while CVP transplanted LV showed lateral wall scarring (red arrows) without obvious ventricular wall thinning. Independently, these observations were confirmed by MRI scans (Figure 4B**, left panel)**. The 4-chamber long-axis view clearly shows the lateral wall thickness (red asterisks). Medium control hearts show a reduction in lateral wall thickness as compared to CVP transplanted hearts. Wall thickness of the hearts were plotted in polar maps according to American Heart Association (AHA) segmentation guidelines^20^. The apical lateral (middle panel, yellow arrow) or mid anterolateral segments were affected due to reduction in blood supply after coronary arteries ligation (Figure 4B**, middle panel)**. Initially, the diastolic wall thickness was similar in the three groups from week 1 to week 4. However, as injury and regeneration proceeded, we observed at 12 weeks a significant (P < 0.05) diastolic wall difference between medium (4.93 mm ± 0.2 mm) and CVP (6.55 mm ± 0.17 mm) transplanted hearts (Figure 4C). There was a large experimental variability at week 12 in the sham group due to the low sample size (n = 3) and this hampered the detection of any significant difference with respect to the CVP group (Figure 4C).

**Figure 4.**
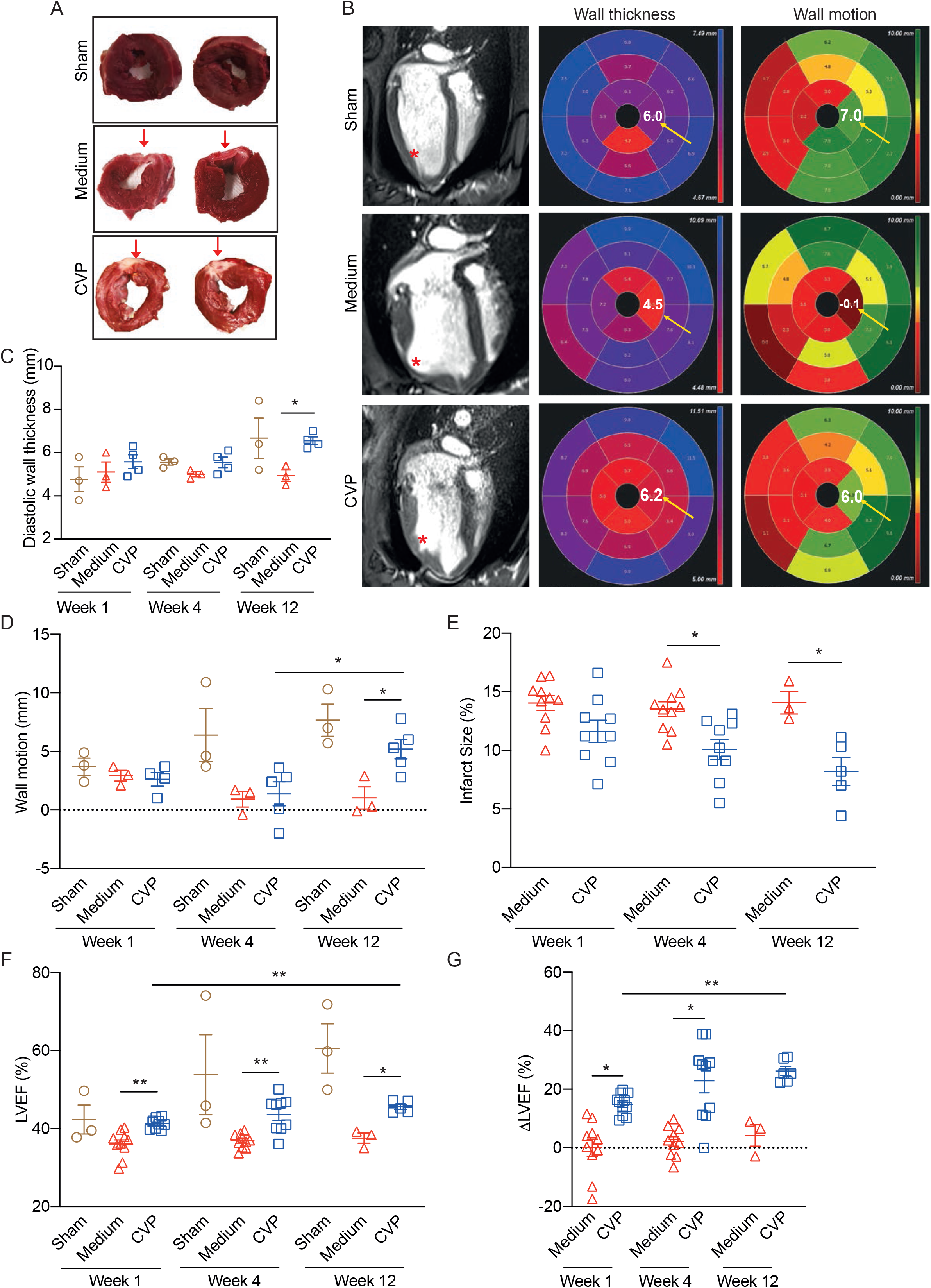
Regenerated infarcted pig heart yields concomitant improvement in heart function. **A)** Representative transverse section of two independent sham, medium and CVP transplanted left ventricles at 12-weeks post transplantation. Transmural infarct region was shown in pinkish white (red arrows) with obvious ventricle remodeling in medium control as compared to CVP transplanted hearts. **B)** Left: 4-chambers long axis MRI slices from sham, medium and CVP transplanted hearts at 12-weeks post transplantation. Significant adverse ventricular remodeling in medium control hearts as compared to sham and CVP transplanted hearts (red asterisks). Center: Representative polar plots (in AHA scale) of left ventricle wall thickness segmentations in sham, medium and CVP transplanted hearts. Functional values in apical lateral segment was highlighted with yellow arrow. Left: Representative polar plots of left ventricle wall motion segmentations in sham, medium and CVP transplanted hearts. Functional values in apical lateral segment was highlighted with yellow arrow. **C)** Quantification of diastolic wall thickness in AHA scale at 1, 4 and 12-weeks post transplantation. Wall thickness in CVP transplanted hearts were significantly thicker than medium control hearts at week 12 (n = 3 - 4). Refer to **(A and B)** for representative images. **D)** Quantification of wall motion at 1, 4 and 12-weeks post-transplantation. CVP transplanted hearts have significantly better wall motion than medium control hearts at week 12 and CVP transplanted hearts at week 4 (n = 3-5). **E)** Infarct size was measured at 1, 4 and 12 weeks post-transplantation. Significant reduction in infarct size from CVP transplanted hearts as compared to medium control at week 4 and 12 (n = 9-10 at weeks 1 and 4, n = 3-5 at week 12). **F)** Left ventricle ejection fraction (LVEF) were determined and measured across 1, 4 and 12 weeks post-transplantation. Significantly higher LVEF in CVP transplanted hearts as compared to medium control hearts at 1, 4 and 12-weeks (n = 9-10, at week 1 and 4, n = 3-5 at week 12). **G)** Change in LVEF (ΔLVEF) from baseline to 1, 4 and 12 weeks are significantly higher (21.35 ± 3.3 %) in CVP transplanted hearts as compared to medium control. Mean ± S.E.M. * P <0.05, ** P < 0.01

Lateral wall motion was also determined by MRI (representative polar plot in Figure 4B, right panel). At week 1, all 3 groups show hypokinetic motion in infarcted segments (2-6 mm). Despite repair and remodeling happening, both medium and CVP groups showed akinetic motion in infarcted segments at 4 weeks (0 - 2 mm) (Figure 4D). However, as regeneration continued to 12 weeks, we observed significant wall motion improvement (P < 0.05) in the CVP hearts (5.2 mm ± 0.8 mm) as compared to medium control (1.033mm ± 0.94mm). Infarcted LV segments improved from akinesis to almost normal motion (> 6 mm). However, medium control hearts remained akinetic. There was also significant improvement (P < 0.05) in CVP transplanted hearts from 4 weeks (1.38 mm ± 1.03 mm) to 12 weeks (5.2 mm ± 0.8 mm) (Figure 4D).

LV infarction size was also determined by MRI measurements (Figure 4E). Initially at 1 week, there was similar infarction size between medium control and CVP transplanted hearts. As the scar develops further to 4 and 12 weeks, there was a significant reduction (P < 0.05) in infarction scar size from 13 – 14 ± 1 % in medium control to 10 ± 0.8 % at 4 weeks and 8 ± 1.2 % at 12 weeks. This strongly suggests a positive regeneration of the infarcted muscle tissue by the transplanted human CVPs.

Collectively, these results suggest that the lateral ventricular wall was adversely affected by the MI with obvious wall thinning, impairment in wall motion and reduction in ejection fraction. However, these changes were significantly restored in hearts transplanted with CVPs at 12 weeks post-transplantation.

The left ventricular ejection fraction (LVEF) of uninjured (sham) pigs are approximately 55 % (Figure 4F). At one week post transplantation, both medium-treated and CVP transplanted hearts had a reduction of LVEF (medium 36 % ± 1.1 %, CVP 42 % ± 3.7 %) as compared to sham. Surprisingly, there was significant improvement between medium- and CVP-treated hearts (P <0.01) as early as at 1 week. This suggests that the transplanted CVPs survived in vivo, proliferated at the injury site, and contributed to the improvement of LVEF. This significant improvement (P < 0.05) was also measured at week 4 (medium 36 % ± 0.6 %, CVP 43 % ± 1.5 %) until the end of 12 weeks (medium 37 % ± 1.3 %, CVP 45 % ± 0.7 %).

The average change in LVEF (ΔLVEF) between medium and CVP transplanted hearts was approximately 21.35 ± 3.3 % as compared to MI control across all the timepoints (Figure 4G). Similarly, there were significant differences (P < 0.01) at week 1 (CVP 15 % ± 1.1 %) and 4 (medium 2.16 % ± 1.62 %, CVP 22.9 % ± 4.1 %) between medium and CVP as well as between week 1 and 12 (26.3 % ± 1.5 %) of CVP transplanted heart (P < 0.01).

Development of tumors is a significant potential concern with the use of pluripotent stem cells in regenerative medicine. During the differentiation on LN221, genes of pluripotency and tumorigenicity were downregulated and we did not observe any signs of tumor formation in our previous study in mice^9^ or in this study in pigs **(Table 1)**. The CT scans at 12 weeks post-transplantation of human CVPs did not yield signs of tumors in the heart or other organs analyzed. Long-term safety is an important aspect for any cellular product to translate into clinical trial. Therefore, we are convinced that the CVPs generated on LN521+221 with our defined, humanized and reproducible protocol are not tumorigenic and that it is likely to be safe as a cellular therapy product.

### Electrophysiological Mapping Reveals Temporary Ventricular Arrhythmia in 50 % of CVP-transplanted Pigs

Accurate discrimination between healthy and infarcted myocardia is crucial to assess the efficacy of CVP treatment. Electromechanical mapping with the 3D-NOGA system simultaneously registers the electrical and mechanical activities of the left ventricle. This enables *in vivo* assessment of myocardial viability. The aim of NOGA was to identify and localize potential arrhythmogenic foci and direct therapeutic procedures.

Seven pigs (n = 1 medium, n = 3 CVP transplanted hearts at 2 weeks, n = 1 each for medium and CVP transplanted hearts at 4 weeks and n = 1 sham pig at 2 weeks) underwent the procedure at 2 and 4 weeks **(Supplementary table 4)**. Unipolar values (mV) provide information on the viability of the tissue by detecting the voltages across the myocardium, while local linear shortening (LLS) values provide information on wall movement. The anterior and lateral wall would be affected when we ligate the LAD and LCX arteries.

The sham pig exhibited a healthy voltage map as shown using unipolar and LLS values in the anterior and lateral walls (Figure 5A). The results were displayed on a LV voltage map (top panels) and bull’s eye polar map (bottom panels). The absence of low voltage segments indicating healthy myocardium (pink areas) are apparent. In contrast, the media control hearts have segments with low voltages and low movement (red area) in the unipolar and LLS map (Figure 5B**, top panels)**. As expected, the bull’s eye polar map demonstrates that the low voltage segments (red areas) and low wall movement (red areas) were at the anterior and lateral walls (Figure 5B**, bottom panels)**. It is noteworthy that overlapping of the low voltage segments and reduced wall movement at the lateral wall is a reliable indicator of scar. The electrophysiological (EP) property of the LV in CVP-transplanted pigs were subsequently determined (Figure 5C). At 2 weeks, we observed similar low unipolar voltages (red areas) and low wall movement (red areas) in the anterior and lateral walls, although to a lesser extent. There was also an overlap of unipolar and LLS segments, indicating the presence of a scar. In parallel to this, the results from the infarct size measurements using MRI measurements from all the pigs (n = 9-10 at weeks 1 and 4, n = 3-5 at week 12) showed a significant reduction in infarct size at 4 and 12 weeks (Figure 4E).

**Figure 5.**
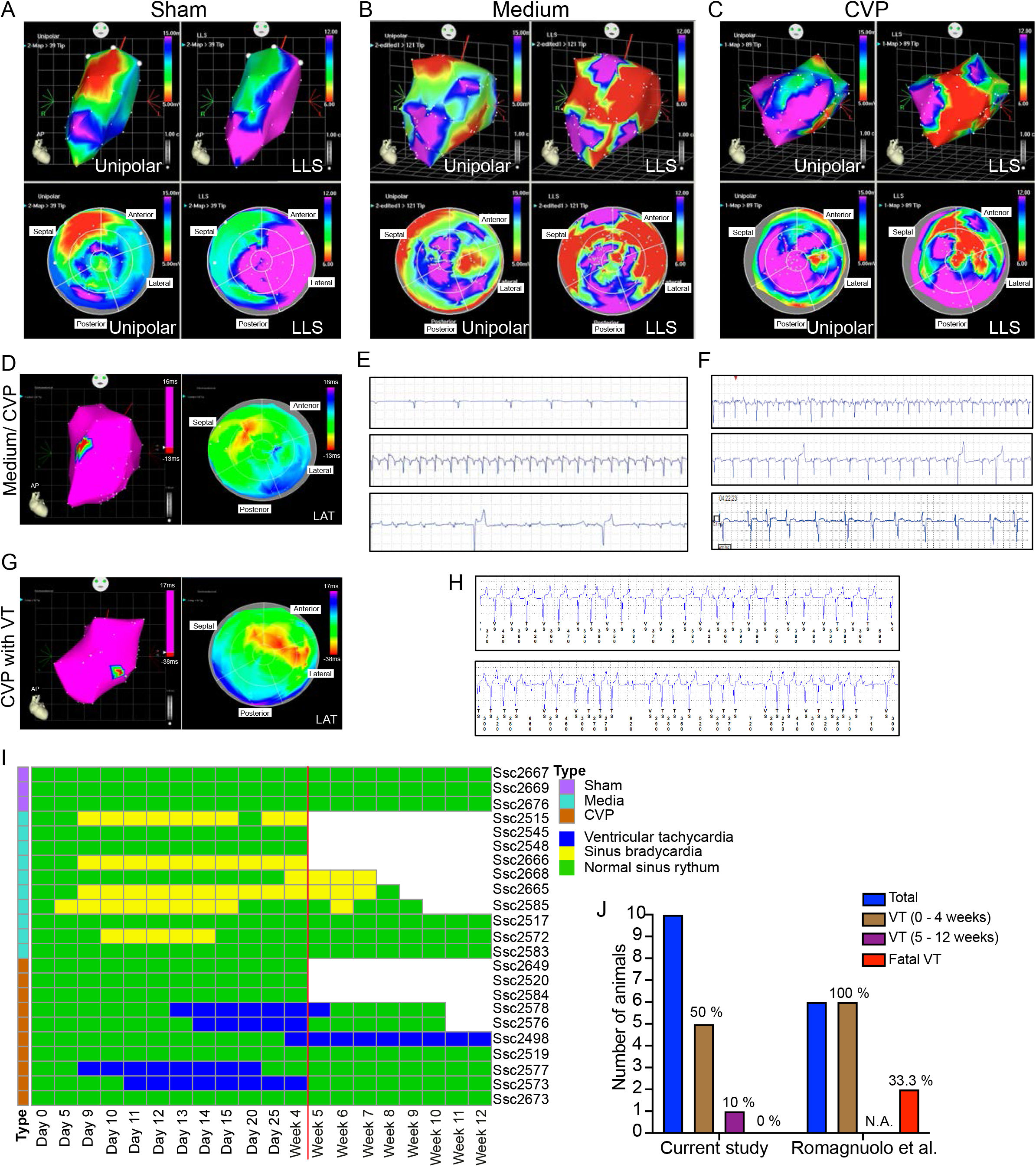
Electrophysiological mapping reveals temporary ventricular arrhythmia in 50 % of CVP-transplanted pigs. Electrophysiology analyses was done on 1 sham pig, 3 medium control pig and 4 CVP transplanted pig at 2 and 4-weeks (Refer to table 1 for more information). **A-C)** Lateral left ventricle endocardial voltage map. **(A)** Healthy uninjured ventricle (sham), **(B)** MI with medium injection. **(C)** MI with CVP transplantation. Unipolar voltage maps indicates the exact positions of infarcted areas and unipolar voltage lower than 5 mV are considered scar (red color), healthy and viable myocardium are indicated as purple. Local linear shortening (LLS) map provides information about wall movement (red = low movement, purple = healthy movement). Results showed overlapping regions with scars (unipolar map) and the non-contractile areas (LLS map). Reduction in scar area and better wall movement in CVP transplanted LV as compared to medium control. **D)** (left) Electro-anatomical maps in medium control or CVP transplanted pigs at 2-weeks post transplantation. It shows the map of a healthy LV without ventricular tachycardia. (right) The local activation time (LAT) map shows LV contraction of a normal electrical activation wave-front where the earliest electrical activation (red areas) is starting in the septum, coming via the left bundle branch and the latest activation (blue areas in the polar map) occurs in the lateral segment. **(E)** Ventricular tachycardia (VT) was observed in two of the CVP transplanted pigs at 2-weeks post transplantation. (left) Electro-anatomical maps in ventricular tachycardia (VT) occurring CVP transplanted pigs at 2-weeks post transplantation. It shows the map of an injured LV with VT. (right) The local activation time (LAT) map shows LV contraction of an abnormal electrical activation wave-front during VT. The earliest activation started at the lateral wall (red areas) instead of septum and ends at the posterior. **F)** Representative ECG traces from implantable cardiac monitor of medium control hearts showing (top) sinus bradycardia, (middle) sinus tachycardia with ST elevation and (bottom) premature ventricular contraction. **G)** Representative ECG traces from implantable cardiac monitor of CVP transplanted hearts (without VT) showing (top) sinus tachycardia, (middle) premature ventricular contraction and (bottom) atrial fibrillation. **H)** Representative ECG traces from implantable cardiac monitor of CVP transplanted hearts (with VT) showing VT. **I)** Heat map showing the overall time course analysis from the implantable cardiac monitor. Sham pigs have normal sinus rhythm throughout the 12 weeks. We observed sinus bradycardia in the 6 out of 10 media controls pigs. For pigs transplanted with CVPs we observed temporary non-fatal VT in 4 out of 10 pigs which resolves itself to sinus rhythm after 5 weeks. One pig (Ssc2498) developed non-fatal VT which started at week 4 and lasted to 12 weeks. **J)** Graph showing the comparisons in this current study and another group that reported the applicability of hESC-derived CMs for MI pig model (Romagnuolo *et al.*). We observed VT in 50 % of our pigs (5 out of 10 pigs) during the first 4 weeks and 10 % (1 out of 10 pigs) of the pigs developed VT at 5 - 12 weeks as compared to 100 % VT occurrence in Romagnuolo’s study (first 4 weeks). No fatal VT was observed in our study as compared to 2 out of 6 pigs that died of fatal VT in Romagnuolo’s study.

To understand electrical connectivity and the heart rhythm, activation mapping was carried out in the pig heart. The medium control pigs showed normal electrical activation (Figure 5D**, left panel)**. The local activation map (Figure 5D**, right panel)** and propagation map **(Supplementary movie 1)** showed earliest electrical activation (red areas) starting from the septum consistent with descending activation from the left bundle branch with the latest activation (blue areas) at the lateral segment as expected. The medium control pigs have normal electrical conductance without ventricular tachyarrhythmia (VT) as confirmed with implantable telemetric ECG recorders (Figure 5E). These traces were analyzed by a blinded clinical electrophysiologist who reported the presence of numerous episodes of sinus bradycardia (top), sinus tachycardia with ST elevation (middle) and premature ventricular contraction (bottom) but no VT.

Analyses of CVP transplanted hearts at 2 weeks showed that the pigs had similar electrical activation as medium control pigs. Initial activation started at the septum and ended at the lateral wall. We also observed conduction into the scar containing human grafts, suggesting electrical conductivity between the host and engrafted cells (Figure 5D**, Supplementary Movie 2)**. Typical ECG traces from the cardiac monitor in CVP transplanted pigs that did not develop VT showed sinus tachycardia (top), sinus rhythm with premature ventricular contractions (middle) and atrial fibrillation (bottom) (Figure 5F).

On the other hand, we observed two CVP transplanted pigs that developed focal VT, as confirmed using both the NOGA machine and implantable cardiac monitor. Earliest activation (red areas) started at the lateral wall and latest activation ended at the posterior segments (blue areas) of the LAT polar map (Figure 5G**, right panel and Supplementary Movie 3)**. Typical VT traces from the cardiac monitor in CVP transplanted pigs that developed VT are shown in Figure 5H. Do note the presence of widened QRS morphology compared to sinus, evidence of atrial ventricular (AV) dissociation and capture beats.

EP mapping was not performed on all the pigs, but all 10 pigs were implanted with telemetric cardiac monitors enabling detection of VT **(Table 1)**. We found out that five out of ten CVP-transplanted pigs developed VT. The VT was temporary and developed at around 10 days post-transplantation and lasted for ∼ 25 days after which the VT resolved itself and the heart returned to normal sinus rhythm (Figure 5I). The complete absence of VT in the remaining five pigs sharply contrasts data obtained in large animals treated with contractile human CMs ^4, 21, 22^ that exhibited extensive arrhythmia and even death. However, we did observe persistent VT in one pig (Ssc2498). VT episodes started at approximately 4 weeks post-transplantation and continued until the end of the experiment without any resolution. The week 1 and 4 MRI scans were not successful because of the high heart rate during VT **(Supplementary table 3)**. Nevertheless, despite persistence of long-lasting episodes of VT, this pig did not die of fatal VT even at end of 12-weeks.

We reviewed the literature on transplanted cells into infarcted pigs and compared the present study against Romagnuolo *et al*.^4^ and plotted a graph to illustrate their differences (Figure 5J). Here, we generated MI and CVP transplantation in 10 pigs. Fifty % of the pigs developed VT during 0 – 4 weeks, and the frequency of VT fell sharply to 10 % during 5-12 weeks, and no death occurred due to VT. In contrast, Romagnuolo *et al*. generated MI and CM transplantation in 6 pigs and 100 % of the pigs developed VT during 0 - 4 weeks. Two pigs died of VT (33.3 % VT-fatality). In summary, results represented here demonstrated that non-beating CVPs are better candidates than mature beating CMs for cell therapy of MI.

## Discussion

The present results support the notion that reproducibly generated non-contractile hESC-derived CVP provide a good starting point for development of cell therapy for MI. Alongside with our previous results obtained in a murine MI model, we confirmed that LN221-drives differentiation of hESCs to CVPs in a highly reproducible and robust manner^9^. This defined and humanized differentiation procedure was documented by multiple time point transcriptome analyses (Figure 1). The CVPs expressed a particular set of genes that were chosen as progenitor marker genes (Figure 1). Day 11 CVPs used for transplantation strongly expressed late CVP progenitor markers such as *ACTC1, ACTN2, ANKRD1, CRHBP, MYH6, TNNI1* and *TNNT2.* Importantly, CVPs differentiated on LN521+221 with different cell lines at 9 or 11 days exhibited high correlation of their expression profiles (R^2^ = 0.97 and 0.96, Figure 1B and 1C) demonstrating the reproducibility of the differentiation method that were transplanted into MI pigs. We have found that differentiating hESCs on human recombinant laminin matrices is highly replicable, e.g. in making endothelial cells (LN521)^23^, photoreceptor progenitors (LN523)^24^, and human keratinocytes (LN211)^25^, highlighting the reliability of the laminin-based differentiation method. Such phenotypic consistency in differentiation is usually not obtained in cells cultured on variable and undefined cellular substrate, such as feeder cells or the mouse tumor extract Matrigel^®^ ^26, 27^ which contains thousands of uncharacterized proteins that vary depending on the tumor batch. We suggest that the use of stable and reproducible differentiation methods is crucial for the development of successful hESC-based cell therapy^2^.

In the pig transplantation study presented here, we used CVPs differentiated on LN221 for eleven days. These cells were still non-contractile, but they expressed more mature CM markers than 9-day cells (Figure 1G - I). Following transplantation into the infarcted region, the cells proliferated and extensively occupied the scar space, building up a new contractile and functional human heart muscle tissue in the host heart. We highlight that 200 million human CVPs were sufficient to improve heart function (at 12 weeks: 1.3-fold improved diastolic wall thickness, 5-fold improved wall motion, 1.7-fold smaller infarct size and 1.2-fold improved LVEF (an overall 21.35 ± 3.3 % improvement)) after permanent ligation of the coronary artery. This represents a significant improvement compared to previous studies, which required 750 million^28^ to one billion contractile CMs in pigs (no improvement in LVEF) and macaques (12.4 % LVEF improvement)^4, 29^. The mechanistic basis behind the effective proliferation of non-contractile CVPs is unclear, but it could possibly be attributed to difference in cell potency between contractile CMs and CVPs, as the former are terminally differentiated while the latter are not^28, 30, 31^.

An important factor in muscle tissue regeneration is access to a sufficient oxygen source, either through existing vessels or neovascularization of the host heart. Here, we showed that despite complete occlusion of the coronary artery, there was rapid appearance of CD31 positive cells and small blood vessels in the regenerating tissue, as well as improved ejection fraction. Similar neovascularization has been reported by others^4, 9, 32–34^. Already at 4 weeks post-transplantation, we observed that the engrafted human grafts generated blood vessels formation. With the possibility of applying cell therapy for repair of heart injury in the future, it may be possible that these CVPs could improve the vasculature of infarcted region.

It is important to note that no signs of tumor growth were observed in this study. In our previous study in mice, transcriptome analyses also showed that the pluripotent hESCs used had their expression of pluripotency and tumorigenic genes quickly reduced after CVP differentiation began^9^. Using the specific combination of LN521 and LN221 as substrate to drive this differentiation, we found that cells from two different cell lines exhibited almost identical properties and similar expression patterns as determined by whole and single-cell RNA-sequencing^35^. Following transplantation, these cells were well engrafted in the immunosuppressed pigs, readily differentiating in the infarcted heart tissue to beating CMs with a rhythm corresponding to that of the host pig hearts.

We report that the transplanted CVPs organized themselves rapidly in the scar, developing mature muscle structures as well as end-to-end junctions between muscle fibers at 4 weeks, similarly to that described by others^4, 6, 29, 34, 36, 37^. The progenitors engrafted and aligned in the scar, expanded and established electrical continuity with the host tissue in contrast to a study by Zhu *et al* that reported the lack of remuscularization after CM transplantation into NHP^1^. Our histological analyses revealed that the cardiac muscle structure at 12 weeks post-transplantation appeared more mature, with clear alignment of muscle fibers beating along the axis of myofibers which developed mature connexin-containing end-to-end junctions. At 12 weeks post-transplantation we observed an overall 21.35 ± 3.3 % improvement of LVEF in CVP transplanted hearts, contrasting with results obtained with contractile CMs. Murry’s group established LVEF improvements of 10.6 ± 0.9 % at 4 weeks and 12.4 % at 12 weeks after CM transplantation in NHP, Laflamme’s group observed no significant improvements in LVEF between medium control and CM transplanted pig hearts^4, 29^.

A similar lack of electrical connectivity has been reported in engineered heart tissue (EHT) constructed from beating CM that have been studied extensively in cardiac regeneration. Querdel *et al.* showed that the transplantation of EHT patches into pigs improved LVEF (24 %) and partial remuscularization of the injured heart. However, electrical coupling needs to be verified in future studies^32^. Bursac’s group reported that EHT patches were able to preserve graft electrical properties but no anterograde or retrograde conduction could be induced between the patch and host CMs, indicating the lack of electrical integration^33^. Similarly, Sawa’s group also reported that induced pluripotent stem cell (iPSC) derived CM sheets could improve cardiac performance in pigs but electrical conductivity was not investigated^38^. Therefore, we went about investigating the electrical conductivity in our human graft. We observed normal sinus rhythm and activation map in 5 of our CVP transplanted pigs. The electrical wavefront was initiated from the septum and went through the low voltage region containing human grafts and ended at the lateral wall **(Supplementary Movie 2)**. This suggests that the human grafts were electrically connected to the host and did not participate in foci activation.

Nonetheless, a critical problem that has surfaced in regenerative cardiology is the development of ventricular arrhythmia in pigs and NHP that have been transplanted with contractile CMs into the infarcted region^4, 29^. In the present study, we observed that 4 of the pigs developed temporary ventricular tachyarrhythmia for about 25 days, after which the heart rhythm returned to normal. Further study is needed to understand the mechanism of VT which we observed in one of our pigs. Could the occurrence of sporadic VT due to the animal, surgery technique or the presence of certain cells in the CVP population showing spontaneous depolarization that leads to focal ventricular tachycardia?

Importantly, we did not observe any death due to the VT episodes and no pigs needed to be euthanized due to VT complications. However, three of the medium controls and two of the CVP-transplanted pigs developed complications (anemia and lung infection, **Supplementary table 3**) probably due to the immunosuppression drugs. The use of improved immunosuppressive drug regiments and other supportive medications will be considered in our future work.

## Acknowledgments

We would like to acknowledge Dr Ru San Tan from National Heart Centre Singapore for his assistance in wall thickness measurements, veterinarians and staffs of SingHealth Experimental Medicine Centre (SEMC) and National University of Singapore (NUS) Comparative Medicine (CM) for their careful and meticulous care for the animals and Duke-NUS Genome Biology Facility for library construction and RNA sequencing. We also thank Dr Paul Lim for his time as a second consultant for the pig ECG traces, technical support team from 10X Genomics and Johnson and Johnson medical electrophysiology team for their support in NOGA mapping.

## Author Contributions

L.Y., L.Y.C, C.T. and M.S. performed the *in vitro* experiments and analyzed the data. L.Y., L.Y.C., C.T., Z.C., M.S., S.J.L. and L.Y. performed the *in vivo* pig experiments and analyzed the data. E.L. analyzed the electrophysiology data from the pigs and N.L. analyzed the CT images of the pigs. S.A. performed the RNA sequencing analysis including 10X spatial transcriptomics. J.G. and E.P. provide technical support and conceptual advice on the bioinformatics. L.Y., S.A., E.P. and K.T. wrote the manuscript. L.Y. and K.T. designed the project, supervised and acquired funding. All co-authors contributed to the preparation of the manuscript.

## Supplementary Figure Legends

**Supplementary figure 1. Characterization of hESCs and CVPs. A)** Representative flow cytometry plots in hESCs using pluripotent markers, Oct3/4 and Tra1-60. **B)** Densitometry analysis of immunoblots known progenitors in Figure 1F. n = 5**. C)** Densitometry analysis of immunoblots CVP signature genes in Figure 1I. n = 5 **D)** Representative flow cytometry plots of H1-derived day 11 CVPs with CVP signature genes (*MYH6, ACTN2, ACTC1, ANKRD1, TNNT2 and TNNI1*).

**Supplementary figure 2. Spatial transcriptome (ST) of pig heart sections engrafted with day 11 human CVPs. A-B)** Spotplots with H&E images and spot overlays depicts the log-transformed normalized expression for pig and human *MYH7, MYL2, TNNC1* and *TPM1* marker genes. The white box shows regions with human CVPs engraftment. **C)** Barplots shows quantified expressions of pig and human gene in the human cells engrafted spots (white box highlighted in panel F-G). T-test, *P*-value < 0.005 *; *P*-value < 0.0005***. Error bars represent standard error of the mean (SEM). **D)** Spotplots with H&E images and spot overlays with unbiased clustering of spots based on global gene expression within individual spots in sham tissue section. E) Pathways enriched in DE genes from across all clusters in sham tissue section (adjusted *P*-values < 0.05). **F)** Functional processes enriched in infarcted and non-infarcted regions of media tissue were computed by *enrichKEGG* (adjusted *P*-values < 0.05).

**Supplementary figure 3. Absence of immune cells in human grafts. A)** A representative H&E image of human graft in infarcted area. **B)** Confocal image of human graft stained with huKu80 (green) and ACTN2 (red) indicating the presence of human cells in the graft. **C)** Confocal images of human grafts stained with immune cell markers CD45, CD20 and CD3, showing the absence of these immune cells.

**Supplementary figure 4. *In vivo* luciferase imaging and H&E staining of CVP transplanted hearts at 12-weeks. A)** Whole heart image demonstrating the presence of the ligation suture (yellow arrows) at the LAD artery after 12-weeks post transplantation confirming the permanent MI model. **B)** *In vivo* IVIS imaging of hearts after CVP transplantation shows the presence of live human cells (bioluminescence signal, red arrows) at 12-weeks post transplantation. **C)** H&E staining on 12-weeks human grafts. Human grafts (indicated as G) are demarcated in dotted yellow lines. Grafts sizes ranges from 1 mm to 10 mm.

**Supplementary figure 5. Confocal images of human grafts. A-B)** Confocal images of human grafts stained with TNNI3 (red), huKu80 (green) and DAPI (blue) at **(A)** 4-weeks and **(B)** 12-weeks post transplantation. **C-D)** Confocal images of human grafts stained with MLC2a (white), huKu80 (green) and DAPI (blue) at **(C)** 4-weeks and **(D)** 12-weeks post transplantation. No atrial-like CMs were present in the infarcted area.

**Supplementary figure 6. Masson Trichrome staining of infarcted region. A-B)** Bright field images of CVP transplanted tissues stained with Masson trichrome at **(A)** 4-weeks post-transplantation and **(B)** 12-weeks post-transplantation. **C-D)** Bright field images of medium control tissues stained with Masson trichrome at (**C)** 4-weeks post-transplantation and **(D)** 12-week post-transplantation. Human graft is demarcated in yellow outline. Collagen deposition (fibrosis) was stained in blue and muscle tissue in purple.

**Supplementary table 1.** Single cell RNA sequencing values (TPM) of the canonical and unique CVP signature genes.

**Supplementary table 2.** Primer sequences

**Supplementary table 3.** Summary of MRI measurements

**Supplementary table 4.** Summary of electrophysiology measurements

**Supplementary Movie 1:** Movie of medium control pig with activation mapping showing scarring in the myocardium with red color

**Supplementary Movie 2:** Movie of electromechanical mappings of CVP transplanted pig without VT.

**Supplementary Movie 3:** Movie of electromechanical mappings of CVP transplanted pig with VT.

