## Supplementary for "PLURIPOTENT STEM CELL-DERIVED CARDIOVASCULAR PROGENITORS DIFFFERENTIATED ON LAMININ 221 REGENERATE AND IMPROVE FUNCTION OF INFARCTED SWINE HEARTS"

Supplementary Figure 1

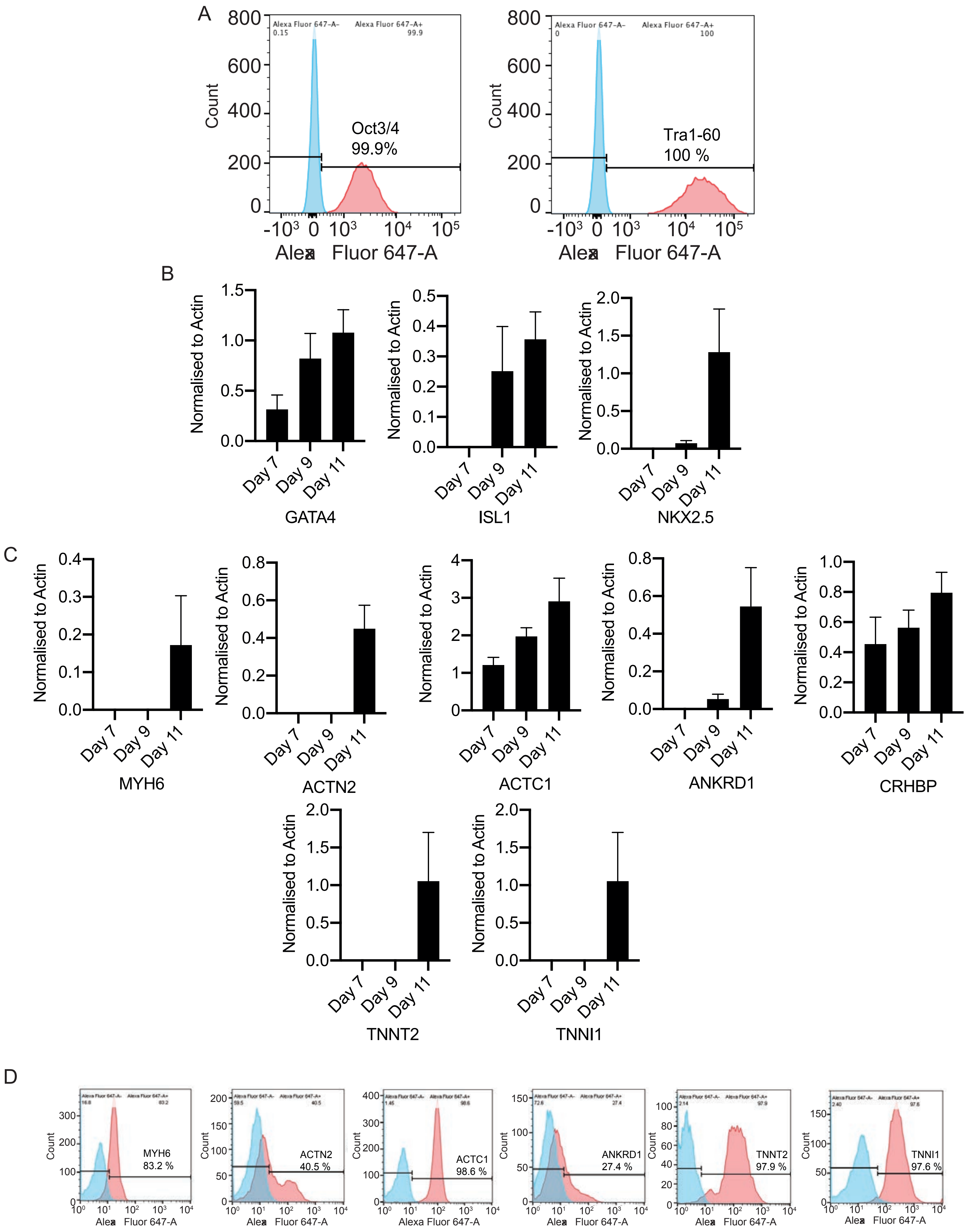

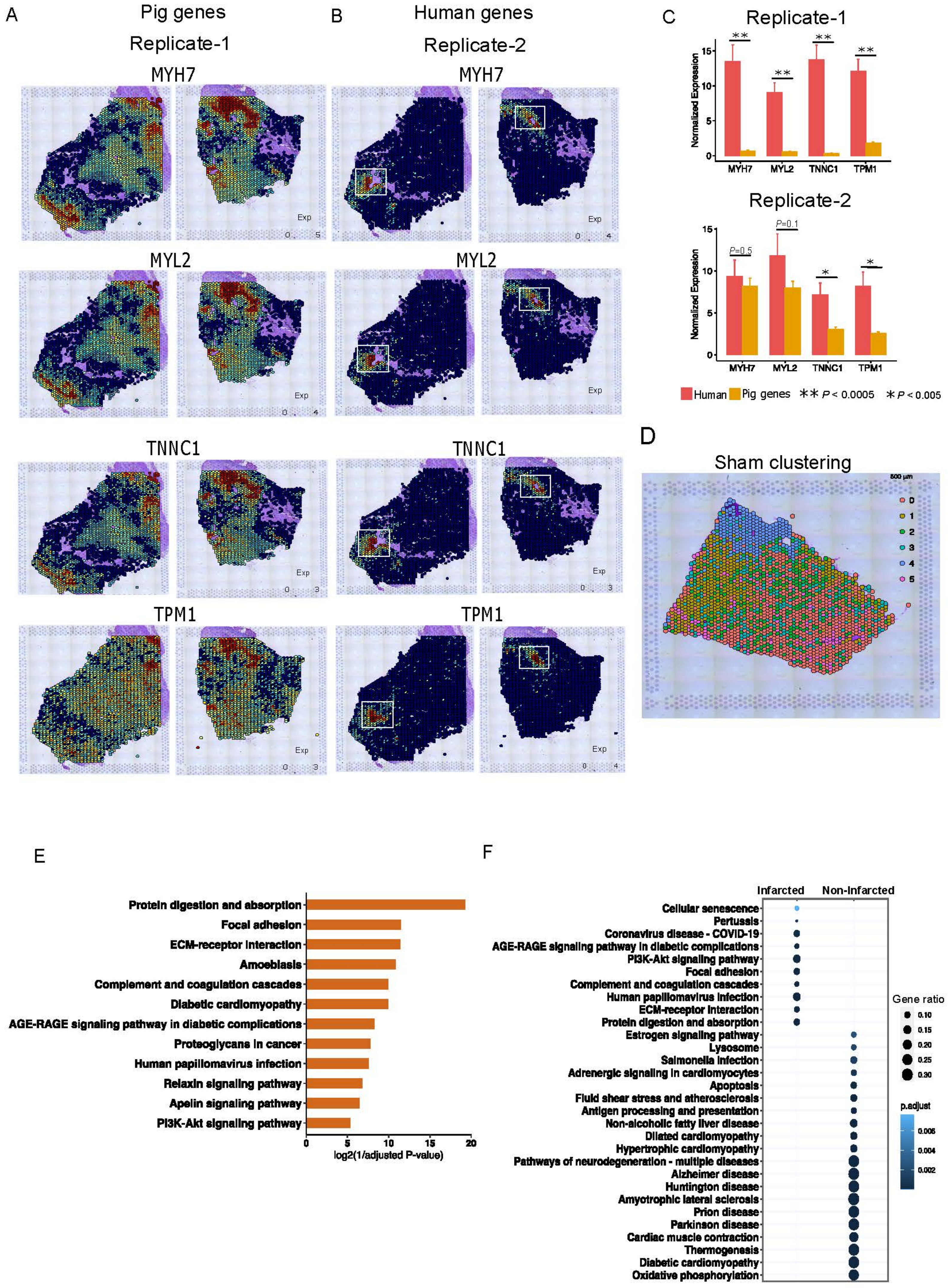
A
Pig genes
Replicate-1
MYH7
MYL2
TNNC1
TPM1
Exp
B
Human genes
Replicate-2
MYH7
MYL2
TNNC1
TPM1
Exp
C
Replicate-1
Normalized Expression
MYH7
MYL2
TNNC1
TPM1
Replicate-2
P=0.5
P=0.1
Human
Pig genes
** P < 0.0005
* P < 0.005
D
Sham clustering
500 µm
0
1
2
3
4
5
E
Protein digestion and absorption
Focal adhesion
ECM-receptor interaction
Amoebiasis
Complement and coagulation cascades
Diabetic cardiomyopathy
AGE-RAGE signaling pathway in diabetic complications
Proteoglycans in cancer
Human papillomavirus infection
Relaxin signaling pathway
Apelin signaling pathway
PI3K-Akt signaling pathway
log2(1/adjusted P-value)
F
Infarcted
Non-Infarcted
Cellular senescence
Pertussis
Coronavirus disease - COVID-19
AGE-RAGE signaling pathway in diabetic complications
PI3K-Akt signaling pathway
Focal adhesion
Complement and coagulation cascades
Human papillomavirus infection
ECM-receptor interaction
Protein digestion and absorption
Estrogen signaling pathway
Lysosome
Salmonella infection
Adrenergic signaling in cardiomyocytes
Apoptosis
Fluid shear stress and atherosclerosis
Antigen processing and presentation
Non-alcoholic fatty liver disease
Dilated cardiomyopathy
Hypertrophic cardiomyopathy
Pathways of neurodegeneration - multiple diseases
Alzheimer disease
Huntington disease
Amyotrophic lateral sclerosis
Prion disease
Parkinson disease
Cardiac muscle contraction
Thermogenesis
Diabetic cardiomyopathy
Oxidative phosphorylation
Gene ratio
p.adjust

Supplementary Figure 3

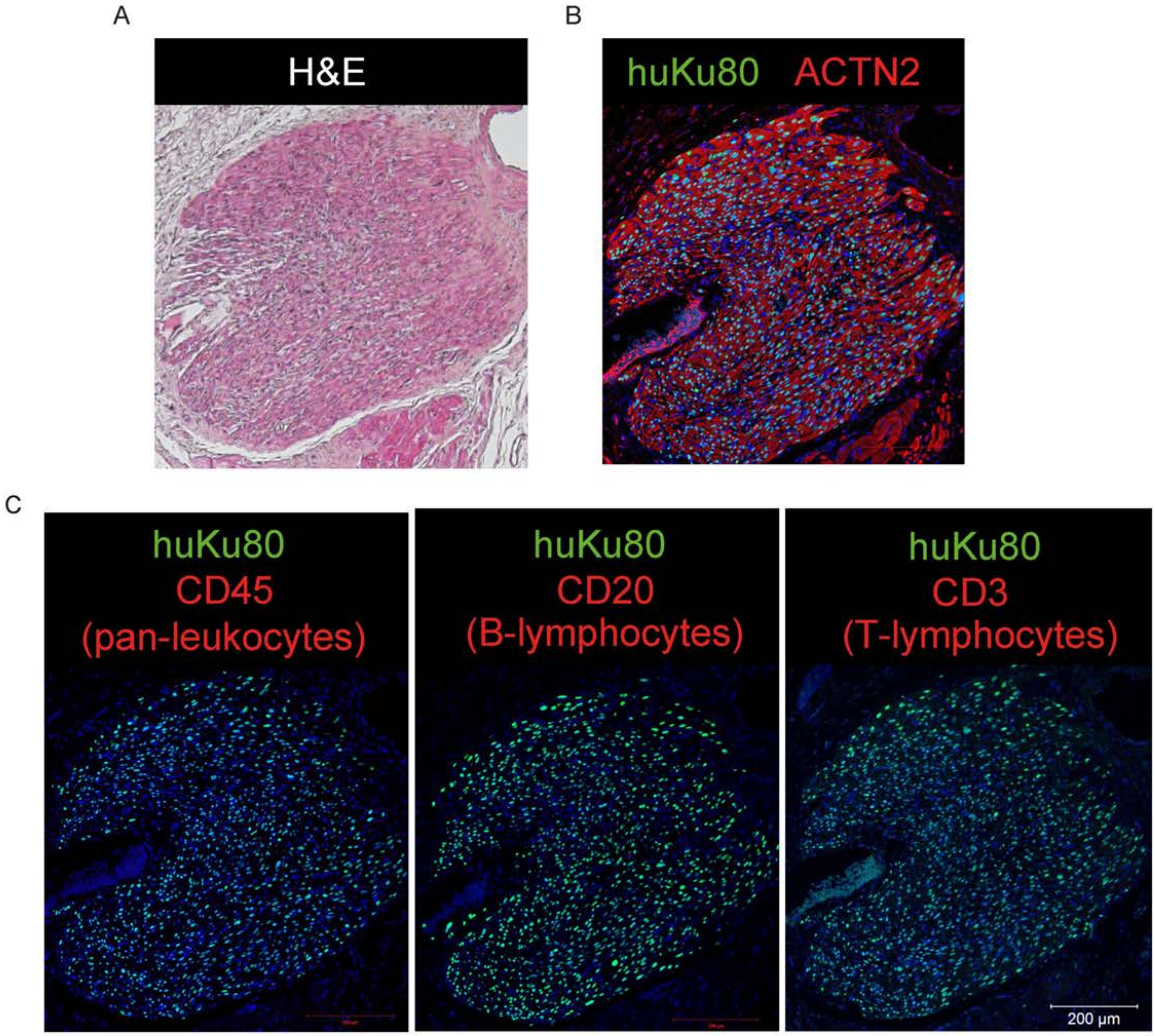

### Supplementary Figure 4

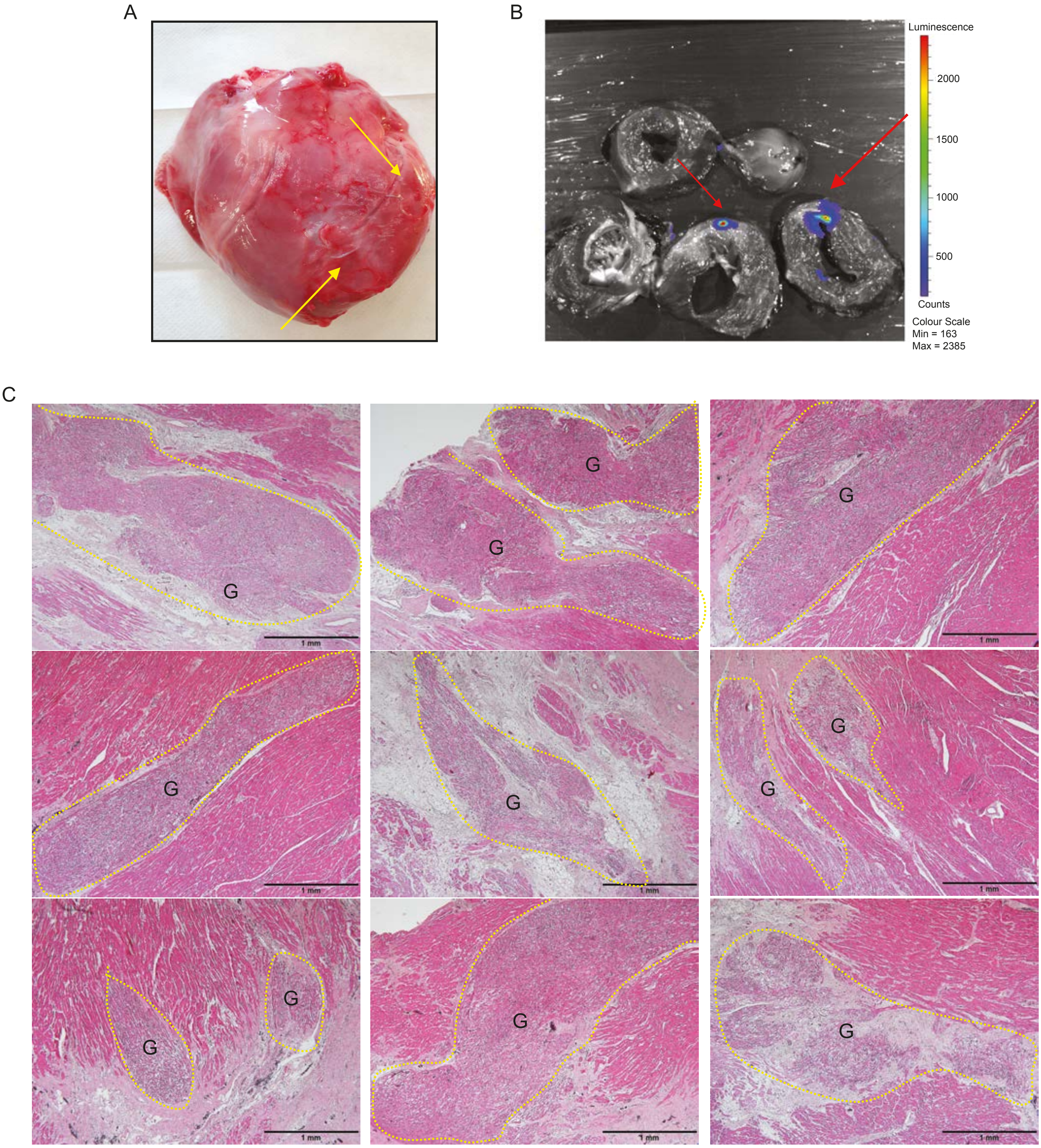

Supplementary Figure 5

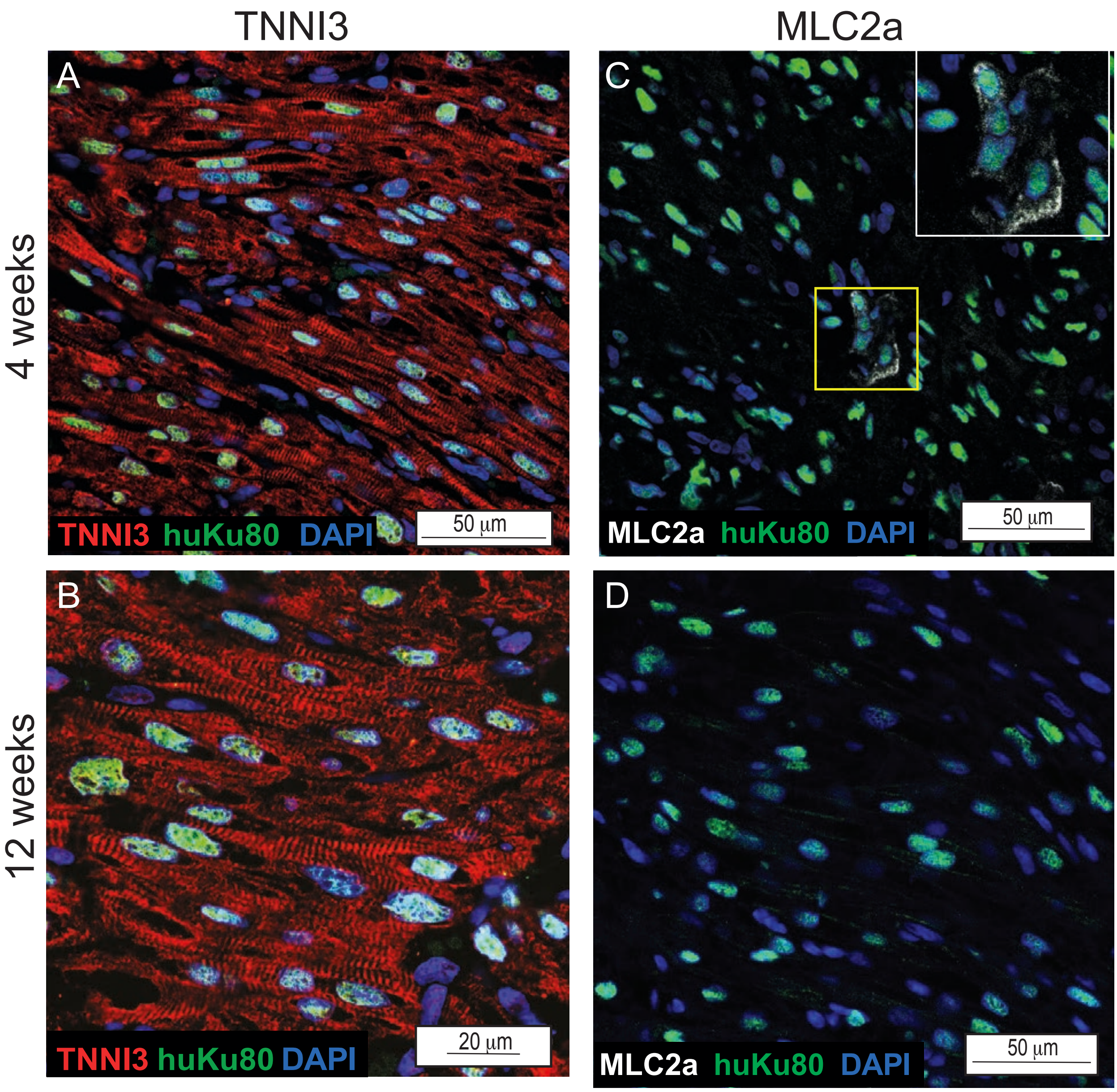

Supplementary Figure 6

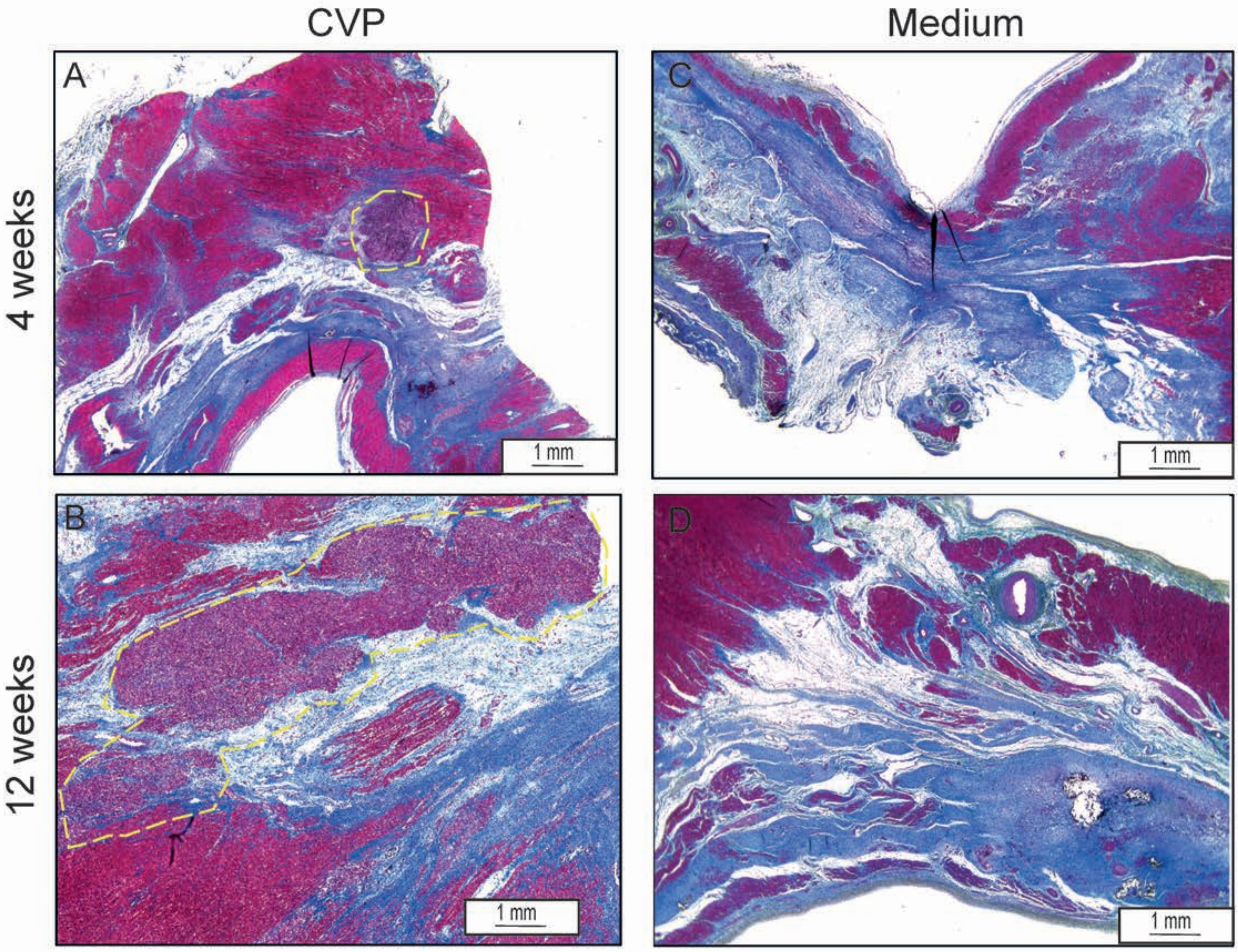

| ID | Gene name | H1_D9 | H1_D11 | H1_Log2FC | H1-padj | HS1001_D9 | HS1001_D11 | HS1001_Log2FC | HS1001-padj |
|---|---|---|---|---|---|---|---|---|---|
| ENSG0000013657 | GATA4 | 33.956 | 32.106 | -0.00801736 | 1 | 35.32333333 | 52.41 | 0.316813635 | 1 |
| ENSG0000001608 | ISL1 | 6.274 | 4.236 | -0.1368087 | 1 | 9.983333333 | 19.66333333 | 1.050546325 | 0.592306025 |
| ENSG0000012805 | KDR | 46.61 | 24.69 | -0.80156639 | 0.1592474 | 26.21333333 | 21.33666667 | -0.377652052 | 1 |
| ENSG0000015740 | KIT | 6.338 | 4.31 | -0.25454001 | 0.9028361 | 6.91 | 6.396666667 | -0.052807565 | 1 |
| ENSG0000016682 | MESP1 | 0.416 | 0.392 | NA | NA | 1.03 | 0.586666667 | NA | NA |
| ENSG0000018307 | NKX2-5 | 3.532 | 30.88 | 3.38455123 | 7.92E-22 | 3.936666667 | 39.9 | 3.231930566 | 4.98E-35 |
| ENSG0000013485 | PDGFRA | 148.302 | 49.024 | -1.0836468 | 0.0341067 | 224.0866667 | 141.8133333 | -0.571467294 | 0.917611945 |
| ENSG0000015925 | ACTC1 | 82.758 | 550.742 | 2.97008319 | 2.50E-13 | 75.32 | 418.5533333 | 2.298648971 | 3.87E-11 |
| ENSG0000007752 | ACTN2 | 2.484 | 28.904 | 3.806661506 | 3.32E-28 | 2.99 | 33.06666667 | 3.70012307 | 1.03E-35 |
| ENSG0000014867 | ANKRD1 | 30.78 | 159.424 | 2.45472823 | 0.0172127 | 63.33 | 149.0133333 | 1.496982184 | 0.003603111 |
| ENSG0000014570 | CRHBP | 2.874 | 61.332 | 4.489434825 | 6.97E-26 | 0.973333333 | 18.94 | 4.118609081 | 1.57E-15 |
| ENSG0000019761 | MYH6 | 15.824 | 470.98 | 5.072899837 | 6.96E-46 | 13.72333333 | 567.3833333 | 5.122266631 | 2.50E-16 |
| ENSG0000015917 | TNNI1 | 36.424 | 210.238 | 2.776113355 | 2.21E-19 | 30.96666667 | 203.17 | 2.609646141 | 8.45E-17 |
| ENSG0000011819 | TNNT2 | 52.404 | 388.866 | 3.054096376 | 5.43E-37 | 50.14 | 356.5433333 | 2.857718916 | 7.43E-16 |

Supplementary Table 2

| Num. | Primer ID | Primer sequence (5' → 3') |
|---|---|---|
| 1 | GATA3-F | TGT CTG CAG CCA GGA GAG C |
| 2 | GATA3-R | ATG CAT CAA ACA ACT GTG GCC A |
| 3 | KDF-F | GGC CCA ATA ATC AGA GTG GCA |
| 4 | KDR-F | CCA GTG TCA TTT CCG ATC ACT TT |
| 5 | PDGFRA-F | TGG CAG TAC CCC ATG TCT GAA |
| 6 | PDGFRA-R | CCA AGA CCG TCA CAA AAA GGC |
| 7 | NKX2.5-F | ACC CTG AGT CCC CTG GAT TT |
| 8 | NKX2.5-R | TCA CTC ATT GCA CGC TGC AT |
| 9 | MESP1-F | TGT ACG CAG AAA CAG CAT CC |
| 10 | MESP1-R | TTG TCC CCT CCA CTC TTC AG |
| 11 | FUT4-F | GAT CTG CGC GTG TTG GAC TA |
| 12 | FUT4-R | GAG GGC GAC TCG AAG TTC AT |
| 13 | KIT-F | CGT TCT GCT CCT ACT GCT TCG |
| 14 | KIT-R | CCC ACG CGG ACT ATT AAG TCT |
| 15 | GATA4-F | GTT GCA CAG ATA GTG ACC CGT |
| 16 | GATA4-R | CGA CAC AAT CTC GAT ATG |
| 17 | TBX5-F | GAG ATA GTC GCT ATC GCC TGG |
| 18 | TBX5-R | AGG TTA TGC TCT CCA ACT ATC |
| 19 | FLRT3-F | CCT CAT CGG GAC TAA AAT TGG G |
| 20 | FLRT3-R | ATG GAT GTC AGA AAG CGA TCA TT |
| 21 | MECR-F | CTG GCG GCC CCT ATC AAT C |
| 22 | MECR-R | AAC ACC TTC GTT CCC TCC AAC |
| 23 | ACTN-F | CAA ACC TGA CCG GGG AAA AAT |
| 24 | ACTN-R | CTG AAT AGC AAA GCG AAG GAT CCC |
| 25 | TNNI-F | CCG GAA GTC GAG AGA AAA CCC |
| 26 | TNNI-R | TCA ATG TCG TAT CGC TCC TCA |
| 27 | MYL4-F | ACT GCC GAC CAG ATT GAA GAG |
| 28 | MYL4-R | CTT GTT GCG GGA AAT GTG CTG |
| 29 | MYH6-F | GCC CTT TGA CAT TGG CAC TG |
| 30 | MYH6-R | GGT TTC AGC AAT GAC CTT GCC |
| 31 | SLC8A1-F | ACA ACA TGC GGC GAT TAA GTC |
| 32 | SLC8A1-R | GCT CTA GCA ATT TTG TCC CCA |
| 33 | ANKRD1-F | AGT AGA GGA ACT GGT CAC TGG |
| 34 | ANKRD1-R | TGT TTC TCG CTT TTC CAC TGT T |
| 35 | ATXN2-F | CTG GGC AGA GGT CGA AAC AG |
| 36 | ATXN2-R | ACA TTT GGA GCC AAC AAC TGA T |
| 37 | CRHBP-F | ATG TCG CCC AAC TTC AAA CTT |
| 38 | CRHBP-R | GCA GGA AAG GAT CGT AGT CCG |
| 39 | ACTC1-F | TCC CAT CGA GCA TGG TAT CAT |
| 40 | ACTC1-R | GGT ACG GCC AGA AGC ATA CA |
| 41 | DLK1-F | CTT TCG GCC ACA GCA CCT AT |
| 42 | DLK1-R | TGT CAT CCT CGC AGA ATC CAT |
| 43 | PLAT-F | AGC GAG CCA AGG TGT TTC AA |
| 44 | PLAT-R | CTT CCC AGC AA TCC TTC GGG |
| 45 | ENO3-F | GGC TGG TTA CCC AGA CAA GG |
| 46 | ENO3-R | TCG TAC TTC CCA TTG CGA TAG AA |

| | | |
|---|---|---|
| 47 | BST2-F | CAC ACT GTG ATC GCC CTA ATG |
| 48 | BST2-R | GTC CGC GAT TCT CAC GCT T |
| 49 | BETA ACTIN-F | GGA CTT CGA GCA AGA GAT GG |
| 50 | BETA ACTIN-F | AGC ACT GTG TTG GCG TAC AG |
| 51 | APOA1-F | CCC TGG GAT CGA GTG AAG GA |
| 52 | APOA1-R | CTG GGA CAC ATA GTC TCT GCC |
| 53 | APOA2-F | CTG TGC TAC TCC TCA CCA TCT |
| 54 | APOA2-R | CTC TCC ACA CAT GGC TCC TTT |
| 55 | GATA6-ASI-F | ACT GTT TGG AGG GAG CGA AG |
| 56 | GATA6-ASI-R | AAG GGC TTC CAC ATC AGT CG |
| 57 | GAPDH-F | AAG GTG AAG GTC GGA GTC AAC |
| 58 | GAPDH-R | GGG GTC ATT GAT GGC AAC AAT A |
| 59 | ISL1-F | GAG GGT TTC TCC GGA TTT GG |
| 60 | ISL1-R | TCC CAT CC TAA CAA AGC ATG T |

Supplementary Table 3

| ID | Cause of death | 1-week LVEDV (ml) | 1-week LVESV (ml) | 1-week LVEF (%) | 1-week Infarct size (%) | 1-week Diastolic wall thickness (mm) | 1-week Systolic wall thickness (mm) | 1-week Wall motion (mm) | 4-week LVEDV (ml) | 4-week LVESV (ml) | 4-week LVEF (%) | 4-week Infarct size (%) | 4-week Diastolic wall thickness (mm) | 4-week Systolic wall thickness (mm) | 4-week Wall motion (mm) | 12-week LVEDV (ml) | 12-week LVESV (ml) | 12-week LVEF (%) | 12-week Infarct size (%) | 12-week Diastolic wall thickness (mm) | 12-week Systolic wall thickness (mm) | 12-week Wall motion (mm) |
|---|---|---|---|---|---|---|---|---|---|---|---|---|---|---|---|---|---|---|---|---|---|---|
| Ssc2667 | | 29.71 | 17.97 | 39.52 | NA | 3.8 | 4.4 | 2.4 | 36.37 | 19.69 | 45.87 | NA | 4.4 | 6 | 4.6 | 48.93 | 19.67 | 59.81 | NA | 5.6 | 11.5 | 5.7 |
| Ssc2669 | | 26.01 | 13.08 | 49.7 | NA | 5.4 | 7.7 | 4.9 | 29.22 | 7.56 | 74.13 | NA | 7.5 | 18.7 | 10.9 | 39.22 | 11.04 | 71.86 | NA | 12 | 20.5 | 10.3 |
| Ssc2676 | | 30.58 | 19.03 | 37.78 | NA | 4.3 | 6.2 | 3.8 | 41.15 | 24.09 | 41.46 | NA | 4.9 | 5.6 | 3.7 | 67.97 | 37 | 45.56 | NA | 6 | 9.9 | 7 |
| Ssc2515 | | 44.68 | 28.7 | 35.77 | 15.1 | NA | NA | NA | 55.88 | 37.06 | 33.69 | 14.9 | | | | | | | | | | |
| Ssc2545 | | 43.4 | 30.48 | 29.76 | 14.2 | NA | NA | NA | 55.78 | 33.7 | 39.59 | 10.5 | | | | | | | | | | |
| Ssc2548 | | 34.89 | 21.01 | 39.78 | 16.3 | NA | NA | NA | 61.87 | 37.86 | 38.81 | 14.5 | | | | | | | | | | |
| Ssc2666 | | 35.22 | 22.48 | 36.17 | 15.1 | NA | NA | NA | 33.29 | 20.97 | 37 | 13.5 | | | | | | | | | | |
| Ssc2668 | Anemia | 32.88 | 20.42 | 37.89 | 12.8 | NA | NA | NA | 35.52 | 23.01 | 35.22 | 13.3 | | | | | | | | | | |
| Ssc2665 | Anemia | 30.59 | 21.02 | 31.29 | 14.5 | NA | NA | NA | 46.77 | 30.41 | 34.99 | 11.6 | | | | | | | | | | |
| Ssc2585 | Anemia | 26.58 | 17.23 | 35.19 | 10 | NA | NA | NA | 65.03 | 40.51 | 37.7 | 11.9 | | | | | | | | | | |
| Ssc2517 | | 37.5 | 22.41 | 40.25 | 14.3 | 4.6 | 4.1 | 3.7 | 51.43 | 32.68 | 36.44 | 13.9 | 4.7 | 3.8 | -0.4 | 57.8 | 37.56 | 35.02 | 15.9 | 6.4 | 5.6 | 0.3 |
| Ssc2572 | | 41.59 | 26 | 37.5 | 11.8 | 6 | 6.3 | 2.1 | 50.13 | 30.89 | 38.39 | 13.7 | 5 | 6 | 1.7 | 41.99 | 25.85 | 38.44 | 12.7 | 4.5 | 4.2 | -0.1 |
| Ssc2583 | | 29.24 | 18.39 | 37.12 | 16.4 | 4.5 | 5.2 | 3 | 62.95 | 39.86 | 36.68 | 17.5 | 5.1 | 4.8 | 1.5 | 88.93 | 53.98 | 39.3 | 13.6 | 6.4 | 6.4 | 2.9 |
| Ssc2469 | | 27.13 | 16.35 | 39.76 | 12.7 | NA | NA | NA | 31.04 | 16.51 | 46.82 | 12.5 | | | | | | | | | | |
| Ssc2520 | | 44.35 | 26.12 | 41.11 | NA | NA | NA | NA | 52.51 | 28.18 | 46.34 | 12.3 | | | | | | | | | | |
| Ssc2584 | | 47.44 | 28.45 | 40.02 | 12.8 | NA | NA | NA | 64.46 | 38.59 | 40.13 | 9.1 | | | | | | | | | | |
| Ssc2578 | lung infection | 45.67 | 25.94 | 43.2 | 16.6 | NA | NA | NA | 66.78 | 40.04 | 40.03 | 13.1 | | | | | | | | | | |
| Ssc2576 | lung infection | 46.33 | 26.86 | 42.01 | 7.1 | NA | NA | NA | 76.5 | 48.91 | 36.07 | 9.1 | | | | | | | | | | |
| Ssc2498 | | 31.96 | 19.35 | 39.45 | 14.3 | Heart rate too fast | | | | | | | | | 0.1 | 65.47 | 37.98 | 41.98 | 8.2 | 5.8 | 6.7 | 5.3 |
| Ssc2519 | | 40.81 | 24.04 | 41.08 | 9 | 5.2 | 7.2 | 3.1 | 49.21 | 26.29 | 46.58 | 10.2 | 5.3 | 6.4 | 2.4 | 91.46 | 48.2 | 47.3 | 10.3 | 7 | 15.4 | 7.8 |
| Ssc2577 | | 44.16 | 25.64 | 41.93 | 11.2 | 5.9 | 8.2 | 1 | 41.27 | 22.23 | 46.14 | 11.7 | 6.1 | 6.7 | -2 | 56.21 | 30.06 | 46.53 | 11.1 | 6.6 | 8.6 | 2.8 |
| Ssc2573 | | 37.29 | 21.73 | 41.72 | 11.2 | 4.9 | 5.7 | 2.7 | 51.64 | 30.47 | 41 | 7.2 | 5 | 7.2 | 3.6 | 72.91 | 41.29 | 43.37 | 7 | 6.4 | 7.1 | 4.1 |
| Ssc2673 | | 27.97 | 15.98 | 42.86 | 9.6 | 6.3 | 6.2 | 3.7 | 50.03 | 24.98 | 50.06 | 5.5 | 5.8 | 5.5 | 2.8 | 57.69 | 31.6 | 45.23 | 4.4 | 7.4 | | 6 |

NA - non applicable

### Supplementary Table 4

| | 2667 (Sham) | | | 2583 (media, 2 wks) | | | 2585 (Media, 2 wks) | | | 2515 (Media, 4 wks) | | | 2573 (CVP, 2 wks) | | | 2577 (CVP, 2 wks) | | | 2584 (CVP, 2 wks) | | | 2519 (CVP, 4 wks) | | | 2519 (CVP, 10 wks) | | |
|---|---|---|---|---|---|---|---|---|---|---|---|---|---|---|---|---|---|---|---|---|---|---|---|---|---|---|---|
| | unipolar | bipolar | LLS | unipolar | bipolar | LLS | unipolar | bipolar | LLS | unipolar | bipolar | LLS | unipolar | bipolar | LLS | unipolar | bipolar | LLS | unipolar | bipolar | LLS | unipolar | bipolar | LLS | unipolar | bipolar | LLS |
| apex | 10 | 2.7 | 13.6 | 6.3 | 3.2 | 8 | 14.9 | 6 | 10.8 | 5 | 2.4 | 10.9 | 21.6 | 6.4 | 7.6 | 13.6 | 4.8 | 2.1 | 10.2 | 4.3 | 9.9 | 7.6 | 4.1 | 12.7 | 7.3 | 5.4 | 18.8 |
| midanterior | 9.4 | 3.4 | 14.2 | 6.1 | 2 | 6.4 | 11.9 | 2.5 | 4.6 | 4.7 | 0.6 | 7.4 | 12.5 | 3.4 | 1.8 | 14.7 | 3.7 | 8.9 | 7.1 | 2.3 | 6.6 | 9.7 | 5.7 | 9.6 | 13.5 | 3.6 | 20.2 |
| anterobasal | 12 | 3.8 | 13.4 | 6.9 | 4.2 | 5.8 | 12.5 | 3 | -8.6 | 11.3 | 5 | 12.6 | 11.1 | 4.3 | 7.6 | 13.7 | 3.5 | 8.5 | 6.8 | 2.2 | 2.3 | 12.2 | 5.7 | 10.4 | 9.5 | 2.8 | 18.3 |
| midlateral | 10.3 | 3.4 | 15.7 | 7.7 | 3.3 | 2.1 | 9.9 | 2.4 | 8.3 | 10.2 | 3.8 | 8.7 | 8.8 | 5 | 10.9 | 10.4 | 2.9 | 5.4 | 5.1 | 3.4 | 5.6 | 11.7 | 4.6 | 11.6 | 5.4 | 1.6 | 6.3 |
| basolateral | 11.6 | 5.9 | 12.6 | 6.7 | 2.8 | 13.9 | 12.8 | 4.2 | 5.8 | 15.3 | 8.2 | 17.2 | 16.5 | 4.1 | 12.3 | 8.9 | 3.3 | 22.9 | 8.5 | 5.3 | 9.5 | 8.4 | 5.1 | 17.2 | 7.3 | 1.8 | 17.3 |
| midposterior | 9.2 | 2.7 | 12.4 | 9.4 | 3.7 | 2.3 | 19.6 | 3.3 | 7.8 | 12.3 | 3.6 | 21.1 | 17.4 | 6.4 | 14.7 | 16.7 | 3.4 | 8.3 | 13.8 | 5.8 | 8.5 | 11.9 | 4.4 | 6.4 | 18.8 | 6.7 | 7.3 |
| posterobasal | 14.4 | 3.8 | 8.6 | 9.1 | 3.7 | 24 | 9.4 | 2.8 | 10.4 | 9.3 | 4.3 | 16.7 | 10 | 3.1 | 14.8 | 9 | 3.6 | 9.8 | 6.8 | 4 | 11.6 | 11.2 | 5.2 | 11.1 | 11.6 | 3.7 | 16.8 |
| midseptal | 12.9 | 1.8 | 12 | 12.4 | 2.8 | 8.8 | 13.9 | 4 | 13.6 | 13.3 | 3 | 13.5 | 14.3 | 5.5 | 11.4 | 14.9 | 5.7 | -1.9 | 11.3 | 3.7 | 11.7 | 12.1 | 4.1 | 12.8 | 19.6 | 4 | 14.6 |
| basoseptal | 5.5 | 1.5 | 9.3 | 9.4 | 5 | -0.5 | 4.6 | 1.8 | 7.9 | 8.4 | 2.1 | 8.7 | 15.4 | 4.1 | 7.7 | 11.5 | 3.2 | 11.4 | 6.1 | 2.8 | 5.3 | 7.9 | 3.7 | 9 | 12.6 | 4.1 | 9.4 |
| Mean | **10.59** | **3.22** | **12.42** | **8.22** | **3.41** | **7.87** | **12.17** | **3.33** | **6.73** | **9.98** | **3.67** | **12.98** | **14.18** | **4.70** | **9.87** | **12.60** | **3.79** | **8.38** | **8.41** | **3.76** | **7.89** | **10.30** | **4.73** | **11.20** | **11.73** | **3.74** | **14.33** |
| Scar segments (bulleye) unipolar Voltage <5mV, bipV <0,5mV | **0** | **0** | | **0** | **0** | | **1** | **0** | | **2** | **0** | | **0** | **0** | | **0** | **0** | | **0** | **0** | | **0** | **0** | | **0** | **0** | |
| %scar unipV/ bipV (9 segments) | **0.00** | **0.00** | | **0.00** | **0.00** | | **11.11** | **0.00** | | **22.22** | **0.00** | | **0.00** | **0.00** | | **0.00** | **0.00** | | **0.00** | **0.00** | | **0.00** | **0.00** | | **0.00** | **0.00** | |
| Segments with normal movement (LLS>6%) | | | 9 | | | 5 | | | 6 | | | 9 | | | 8 | | | 6 | | | 6 | | | 9 | | | 9 |
| hypokinetic segments (2-6% LLS) | | | 0 | | | 3 | | | 2 | | | 0 | | | 0 | | | 2 | | | 3 | | | 0 | | | 0 |
| diskinetic segments(- %) | | | 0 | | | 1 | | | 1 | | | 0 | | | 0 | | | 1 | | | 0 | | | 0 | | | 0 |
| akinetic segments (0-2% LLS) | | | 0 | | | 0 | | | 0 | | | 0 | | | 1 | | | 0 | | | 0 | | | 0 | | | 0 |
