## Extended Material and Methods for "PLURIPOTENT STEM CELL-DERIVED CARDIOVASCULAR PROGENITORS DIFFFERENTIATED ON LAMININ 221 REGENERATE AND IMPROVE FUNCTION OF INFARCTED SWINE HEARTS"

**Maintenance of human embryonic stem cells (hESCs)**

All human pluripotent stem cell studies were carried out in accordance with approval from National University of Singapore's Institutional Review Board (IRB 12-451). All pluripotent hESCs H1 (WiCell Research Institute, WA01) and HS1001[1] were maintained on 6-well culture plates coated overnight (4°C) with 10 mg/ml of recombinant LN-521 (Biolamina AB, LN521) in PBS. Maintenance medium Nutristem® (Biological Industries, 05-100-1A) was refreshed daily[1]. Routine monitoring of pluripotent markers (POU5F1 (Santa Cruz, sc-5279) and Tra1-60 (Millipore, MAB4360)) by flow cytometry and genomic stability by karyotyping were performed. Cells were split at 100,000 cells per well and were passaged at 80 % confluence by gentle dissociation with TrypLE (ThermoFisher, 12563011) at 37 °C for 8 mins followed by pipetting to dissociate the single cells. The cell suspension was then collected and centrifuged at 800 rpm for 4 mins. Supernatants were discarded and the cell pellets were resuspended in 1mL of warm Nutristem® medium. Bright field images were taken with Leica microscope.

**Cardiovascular progenitor differentiation protocol**

Pluripotent cells were seeded at 6 million cells into 10-cm$^2$ dishes (ThermoFisher, 150464) coated with combination matrices of (3.33 μg/ml) LN-521 and (10 μg/ml) LN-221 on day 0 in 5ml of PBS. This protocol was based on our previous study[2]. At confluence (day 4), the cells were exposed to differentiation medium (RPMI 1640 (ThermoFisher, 11875-093) with B27 supplement without insulin (ThermoFisher, A1895601) and 10 μM of CHIR99021 (Tocris, 4423) for 24 hours. The next day (day 5), the medium was removed and replaced with differentiation medium. At day 7, the medium was changed to differentiation medium with the addition of 5 μM of IWP2

(Tocris, 3533) for 2 days. Wells were replaced with differentiation medium at day 9 and day 11. A 10-cm$^2$ dish can generate ~ 20 million cells. Therefore, in order to achieve 200 million cells for 1 pig, we performed differentiation in ten 10-cm$^2$ dish. Schematic presentation of the differentiation protocol is shown in figure 1A.

**Generation of bulk RNAseq library**

Bulk RNA sequencing data was generated in our previous publication[2]. Briefly, human embryonic stem cells from H1 and HS1001 cell lines were cultured in wells coated with LN-521+221 at different days from the differential protocol (as described in Yap *et al.* 2019[2]). RNA was extracted from triplicates at day 0, 1, 3, 11, 14, 20, 30 and 90 and from 5 replicates at days 5, 7 and 9. Libraries were constructed with TruSeq stranded total RNA library Prep Kit. The sequencing data was deposited in NCBI's Gene Expression Omnibus Omnibus[3] and are accessible through GEO Series accession number GSE100725.

**Bulk RNA-sequencing data processing**

To identify the unique CVP progenitor signature genes that specifically mark days 9 and 11, the genome wide transcripts per million counts were extracted from our earlier study[2] [GSE100725]. To account for reproducibility across the H1 and HS1001 cell lines we performed correlation analysis independently at days 9 and 11. The Spearman's ranked correlation was computed and plotted using *heatscatter* function from LSD R package (v4.1-0). The selected CVP markers genes that are significantly differentially expressed in two cell lines and those clearly distinguishes day 9 vs day 11 compared to other timepoints expression were extracted. These marker genes expression heatmaps were generated using ComplexHeatmap R Bioconductor package (v2.6.2).

**Quantitative Real-Time Polymerase Chain Reaction (qPCR)**

Total RNA from Day 7, Day 9 and Day 11 was isolated using Total RNA Purification Kit (Norgen Biotek, 17200) and quantified on the NanoDrop 1000 Spectrophotometer (Thermo Fisher Scientific). Complementary DNA (cDNA) synthesis from 5 μg of total RNA was performed using iSCRIPT™ Reverse Transcription Supermix (BIO-RAD, 1708841) and T100 Thermal Cycler (Bio-Rad). Quantitative Real-Time PCRs (qPCR) were performed with CFX384 Real-Time PCR system (BIO-RAD), using a total volume per reaction of 10 μl containing 1X SYBR™ Select Master Mix for CFX (Applied Biosystems™, 4472942), 2.5 μl of 1 μg cDNA template, 1 μl of 10 μM forward and reverse primers mix and 1.5 μl sterile RNase-free water. The following thermal profile was applied: 1 cycle at 95 °C for 3 min, 40 cycles at 95 °C for 10 sec and 60 °C for 1 min, melt curve cycle from 65 °C to 95 °C for 2.5 min. Direct detection of PCR products was monitored by measuring the fluorescence produced due to SYBR Green dye binding to dsDNA after every cycle. The nucleotide sequences of forward and reverse primers for *MECR, APOA1, APOA2, BST2, ENO3, PLAT2, GATA6, GATA3, KDR, PDGFRA, FLRT3, ACTN2, TNNT1, MYH6, SL8A1, ANKRD1/CARP, CRHBP, ACTC1, DLK1, NKX2.5, ISL1, MYL4, MESP1, FUT4, KIT, GATA4, TBX5, ATXN2, GADPH* and *ACTB* were tabulated in Supplementary Table 2. Cq values were normalized to MECR and results plotted as relative expression units.

**Western Blot**

Cells were dissociated with TrypLE™ Select Enzyme (Gibco, 12563011), centrifuged into pellet in a microcentrifuge tube and kept at -80$^{o}$C until all time points (Day 7, 9 and 11) were collected. Cell pellets were washed once in ice-cold PBS and total protein was harvested using ice-cold RIPA Lysis and Extraction Buffer (Thermo Fisher Scientific, 89900) supplemented with 1X Halt™ Protease Inhibitors Cocktail (Thermo Fisher Scientific, 78429). Total protein was quantified using the BCA assay (Thermo Fisher Scientific, 89900). 10 μg total protein was denatured at 100 $^{o}$C for 5 min with 1X Laemlli and RIPA supplemented protease inhibitor. Total protein was run on 4-12% Bis-Tris NuPage Gel (Invitrogen, NP0323BOX) under reducing conditions at 120 V for 90 minutes, protein was subsequently transferred to methanol-activated PVDF membrane at 75 V for 90 mins. Membranes were blocked in 5 % (w/v) skim milk in Tris-buffered saline with Tween-20 (TBST), after which membranes were stained for either MECR (Thermo Fisher Scientific, PA5-54555, 1:250), GATA4 (Abcam, ab124265, 1:1000), ISL1 (Abcam, 86472, 1:1000), NKX2.5 (Santa Cruz Biotechnology, sc-14033, 1:250), MYH6 (Abcam, ab50967, 1:1000), ACTN2 (Sigma, A7811, 1:1000), ACTC1 (Sigma, SAB5600071, 1:10000), ANKRD1/CARP (Millipore, MABS1228, 1:250), TNNT2 (Abcam, ab91605, 1:5000), TNNI1 (Sigma, AV42117, 1:1000) or actin (Millipore, MAB1501R, 1:10000). Membranes were further washed with TBST and incubated with horseradish peroxidase-conjugated secondary antibodies in 5 % (w/v) skim milk in TBST. Membranes were finally washed and developed using Amersham ECL Prime Western Blotting Detection Reagent (Cytiva, RPN2232) and detection was performed using a ChemiDoc MP Imaging System (Bio-Rad). Images are representative of 5 independent differentiation batches.

Densitometry performed using ImageJ and normalized to actin. Data represents average of 5 independent differentiation batches.

**Flow Cytometry**

Specific markers were probed with antibodies using flow cytometry. Cells were dissociated and centrifuged into pellet in a microcentrifuge tube. The cells washed with 1X DPBS, then fixed with reagent A of the Fix and Perm kit (ThermoFisher, GAS004) for 15 mins at room temperature, followed by a wash with 1 mL of 1% (w/v) BSA (Sigma, A7906) in PBS and spun at 14,000 rpm for 1 minute. Subsequently, cells were permeabilized with reagent B in the Fix and Perm kit and incubated together with primary antibodies: POU5F1 (Santa Cruz, sc-5279, 1:20), Tra1-60 (Millipore, MAB4360, 1:50), TNNT2 (Abcam, ab91605, 1:100), ISL1 (DSHB, 39.4D5, 1:20), NKX2.5 (Santa Cruz Biotechnology, sc-14033, 1:50) or appropriate isotype control, for 15 mins at room temperature. After incubation, the cells were washed with 1% (w/v) BSA in PBS, spun and incubated in the dark for 15 mins with the appropriate Alexa-conjugated secondary antibody (ThermoFisher) at 1:1000 dilutions in 1% (w/v) BSA in PBS. Finally, the cells were washed and resuspended in 250 μL of 1% (w/v) BSA in PBS prior to analysis in FACS Fortessa (Becton Dickinson). Data were analyzed using FlowJo software (Version 8). Cell gating (% expression) was done at the intersection between the isotype control and the marker expression[4]. There were 5 replicates in each group.

**Generation of 10X Visium Spatial Transcriptomics**

One week following surgery, we euthanized the animals and sectioned the left ventricle into 5 cross-sectioned rings. The rings were imaged under the IVIS machine and tissue with positive luciferase were sectioned into smaller pieces. The pieces (7 mm x 7 mm) were immediately snapped freeze in liquid nitrogen and sectioned into 10 micron thickness onto blank slides. To confirm the location of the human cells, we performed H&E and Ku-80 staining. Once the location was found, the next subsequent section was placed onto the 10X gene expression slide for downstream processing in accordance to 10X Visium Spatial Gene Expression user guide (https://support.10xgenomics.com/spatial-gene-expression/library-prep/doc/user-guide-visium-spatial-gene-expression-reagent-kits-user-guide).

We achieved a tissue coverage area of ~ 60% [(0.6 % x 5 000 total spots) x 50,000 read pairs/spot = 150 million total read pairs] and RNA were sequenced using Novoseq 6000 system (150 paired-end reads).

**Spatial Transcriptomics data processing**

To segregate the spatial transcriptomics (ST) reads belonging to the transplanted human CVPs from the host pig heart reads, we prepared a combined genome reference from human and pig ensembl genomes (GRCh38 and Ssus11, v102) using *mkref* from 10X spaceranger (v1.2.1) [https://support.10xgenomics.com/spatial-gene-expression/software/overview/welcome]. The raw reads were aligned to combined reference genome and the summarized UMI counts for each spot on the visium spatial transcriptomics array were generated using spaceranger *counts*. The mRNA count matrices were generated by adding intronic and exonic reads for each gene in each location. The paired histology H&E images were processed using spaceranger to select locations covered by tissue by aligning to spot locations with fiducial border spots in the histology image. This enabled to overlay the quantified counts for genes in each spot. Only the spots under the tissue sections were retained for the downstream analyses. Raw counts, images, spot-image coordinates and scale factors were imported into STutility R package (v0.1.0)[5]. The resultant gene-spot matrix generated was analysed with the STutility and Seurat (v4.0.1) packages. As a quality control (QC) we removed genes that are expressed in less than 1% of spots and less than 1% of total reads. Spots expressing extremely low or high number of genes were removed (below 5th or above 95th percentile). Spots with more than 10% of their total gene count coming from mitochondrial and ribosomal genes were also discarded. The QC step resulted in 11328 genes and 9269 spots (2187, 2087, 2948 and 2047 spots for sham, medium, replicate-1 and replicate-2 respectively).

The post-QC gene-spot matrix counts were normalized across spots using regularised negative binomial regression (*SCTransform* function from Seurat package). Normalization across spots was performed with by regression of number of genes per spot. Dimensionality reduction was performed using non-negative matrix factorization (*RunNMF* function in STUtility) for each tissue section. The number of factors/dimensions for the factor analysis were empirically set based on the spatial patterns in each tissue (9 and 16 factors for sham and medium tissues, and 25 factors each for replicate tissues). Spots in each tissue were clustering using the Shared Nearest Neighbor (SNN) algorithm (using *FindNeighbors* and *FindClusters* functions from Seurat package) at resolution 0.4 each for sham, medium and replicate-2, and a resolution 0.3 for replicate-1. Identified clusters in each section were visualized in spatial context by overlaying spot over H&E images (with spot size scaling) using *ST.FeaturePlot* function from STutility (Figure 2J-L). To identify markers within each cluster, we compared each cluster versus all others using the *FindAllMarkers* function (min.pct = 0.1, adjusted *P*-values <0.05 non-parametric Wilcoxon rank sum test). The functional analysis for differentially expressed genes in sham and medium sections was performed using *enrichKEGG* from ClusterProfiler R package (v3.18.1)[6].

We extracted the clusters with human CVPs engraftment from replicate-1 and replicate-2 by computing the number of human genes expressed in cluster (tables in Figure 2J-L). The expression of human and pig marker genes were plotted using *FeatureOverlay* function in STUtility (Figure 2F-G). We quantified and compared the expression levels of pig and human CVP markers in all the spots from the engrafted clusters. The statistical differences across the pig and human CVP markers was computed using *t.test* R function (Figure 2H). To account for reproducibility across the replicates, we computed the Spearman's ranked correlation of pig and human genes. The correlation across the replicates was visualized using *pairs* R function (Figure 2I). The common genes expressed in human CVPs engrafted clusters from replicate-1 and -2 were extracted and plotted with *EnhancedVolcano* R package (v1.8.0) (Figure 2M). Functional enrichment analysis was performed on the common genes using *gseKEGG* from *clusterProfiler* package[6]. The enriched pathways with adjusted *P*-value below 0.05 were plotted using *ggplot2* R package (v3.3.3) (Figure 2N).

**Immunosuppressed Pig Model**

Either gender of pigs (Sus scrofa) at 3 months old, 13-15 kg were purchased from SingHealth Experimental Medicine Centre (SEMC, Singapore) and used for all experiments. Immunosuppression was given to prevent rejection of the human cells in the pig heart. Five days before the surgery, cyclosporine (Novartis, ADP835296) was given in the diet twice daily at 15 mg/kg and maintained throughout the experiment. First dose of Orencia® (Bristol-Myers Squibb, ABT4318) at 12.5 mg/kg was given on the day of surgery via intravenous injection and once every 2 weeks until the end of the experiment. Corticosteroid immunosuppression, methylprednisolone (*Vem ìlaç*, 011001) was injected via intraperitoneally once daily (250mg on surgery day, 125 mg for first 2 weeks after transplant and 62.5 mg until the end of the experiment). On the first 14 days post-surgery, antibiotics (Betamox (every other day) and Baytril (once daily)) were administrated to prevent infections. Analgesic (buprenorphine (twice daily)) were also administrated on the day of surgery and after electrophysiology mapping. To overcome the side effects of immunosuppression, iron capsules (26mg, Now Foods) were given through diet every other day and ranitidine (twice daily) to prevent stomach ulcers. All the procedures involving animal handling were performed with prior approval and in accordance with the protocols and guidelines of SingHealth's Institutional Animal Care and Use Committee (IACUC) (2018/SHS/1426).

**Generation of myocardial infraction model and CVP transplantation**

To track the biodistribution and viability of the progenitors, we employed our previously generated luciferase-labelled H1 cells in the pig experiments[2]. Sus scrofa were anesthetized, intubated, and maintained through a ventilator. A limited left lateral thoracotomy was performed to expose the heart. The first branch of left coronary artery (D1) and first branch of left circumflex (M1) were permanently ligated with nonabsorbable suture (B Braun, C0026003). This consistently gave a scar size of approximately 15 % of the left ventricular anterior wall[7]. In treated group, single cell CVPs (200 million cells) in RMPI medium at 1 ml was intramyocardially injected into infarcted and peri-infarcted myocardium at several points (~ 10 sites). For the medium control pigs, 1 ml of RPMI medium was similarly intramyocardially injected into the infarcted region. Chest was closed-up and the animal was monitored closely after the surgery. Implantable loop recorder was implanted into the chest wall for daily ECG recording and transmission to Reveal LINQ$^{TM}$ cardiac monitoring system (Medtronic).

Animals were maintained for 4 or 12 weeks. Schematic representation of the surgery procedure is shown in figure 2A.

**Bioluminescent Measurements**

After euthanisation of the pig, D-luciferin (15mg/ml in 2ml volume) (Perkin Elmer, 122799-5) were administered into the excised whole heart via injection into both coronary arteries. Ten minutes after injection of D-luciferin, the right ventricle was removed and left ventricle was cross sectioned into 5 rings. All 5 rings were placed on the IVIS Spectrum imaging platform (Perkin Elmer) and 2D bioluminescent image were taken to calculate bioluminescent signals and total photons emitted from the heart area. Areas on the tissue rings with positive signals were sectioned and fixed in 4 % PFA (Sigma, 28908) for downstream tissue processing and immunostaining.

**Immunohistology staining**

Pig hearts were coronally dissected and transversely sectioned into five slices (15 mm thick) at weeks 4 or 12 post MI. The infarcted area and border zone tissues were further dissected and fixed in 10 % neutral buffered formalin (Sigma, HT501128) overnight at room temperature, processed and paraffin-embedded for histological analyses. Tissues were sectioned into 5 $\mu$m thick sections followed by deparaffinization and heat induced antigen retrieval using sodium citrate buffer, pH 6 (Sigma, C9999) for 10 mins. After which, sections were washed thrice with 1 X PBS for 5 mins each and incubated with goat serum (Sigma, G9023) and 0.2 % Triton X-100 (Sigma, T8787) for 30 mins. Sections were then incubated with specific primary antibody: anti-TNNT2 (Abcam, ab91605,1:100), anti-Ku80 (Cell Signaling, 2180S, 1:300), anti-Ku80 conjugated 488 (Abcam, ab198586, 1:100), anti-MLC2v (Abcam, ab79935, dilution?), anti-ACTN2 (Sigma, A2172, 1:500), anti-N-cadherin (Sigma, C3678, 1:100), anti-CX43 (Sigma, 6219C, 1:200), anti-CD31 (Abcam, ab28364, 1:100), anti-TNNI3 (Novus, NBP1-56641, 1:100), anti-MLC2a (Sigma, HPA013331, 1:100), anti-Ki67 (Abcam, ab15580, 1:100), anti-PPH3 (Cell Signaling,9701,1:100), anti-CD45 (Bio-Rad, MCA1447, 1:100), anti-CD20 (Biocare Medical, ACR3004B, 1:100) and anti-CD3 (DAKO, A0452, 1:100) 4 °C overnight. Next day, sections were washed thrice with 1 X PBS for 5 mins each. Alexa-conjugated secondary antibody (ThermoFisher, 1:1000) and DAPI (ThermoFisher, D1306, 1:5000) were added for 1 hr and washed thrice with 1 X PBS for 5 mins each. To suppress autofluorescence, the sections were incubated with Sudan Black (Sigma, 199664) for 20 mins, washed thrice with 1 X PBS and then mounted with ProLong Gold antifade mountant (ThermoFisher, P36930). Slides were examined using LSM 710 Carl Zeiss confocal microscope.

Proliferation rate of the engrafted cardiomyocytes was determined from sections stained for Ki67 and PPH3. Analysis were done by manually counting of the Ki67+ and PPH3+ nuclei from the average of five fields per heart (40 X objective) by a blinded researcher. The proliferation rate was expressed as % of proliferation = (number of Ki67+/PPH3+) divided by the (number of Ku80+ human nuclei).

**Hematoxylin and Eosin Staining**

Sections were deparaffinized in 2 changes of Histoclear (Nanoserv, 64110-04), 2 mins each, followed by 100% ethanol and 70% ethanol, washing in water for 5 mins. After rehydration, sections were incubated in Hematoxylin (Biomed Diagnostics, 3801520) for 10 mins, washed thrice in water for 5 min each followed by 1 min incubation in Eosin (Leica, 3801601). For dehydration, slides went through series changes of ethanol (70% to 100% ethanol), 3 changes of Histoclear and mounted with mounting media (Leica, 1407093626). Slides were examined using Leica DMi8 microscope.

**Masson Trichrome Staining**

Deparaffinized and rehydrated sections were fixed in Bouin’s (Sigma, HT10132) solution for 1hr at 60°C. Rinsed sections in running tap water for 15 min to remove the yellow color. Sections were stained in Weigert’s Iron Hematoxylin solution (Merk, 1.15973.0002) for 10 min, washed in water for 5 mins, stained in Biebrich Scarlet-Acid Fuchsin solution (Sigma, B6008 and F8125) for 3 min and rinsed with water. After that, differentiate in phosphomolbdic-phosphotungstic acid solution (Nanoserv, 19400 and 19500) for 10 min, transferred sections directly to methylene blue solution (Merk, 1.15943.0025) and stained for 10mins. Rinsed briefly in water and differentiated in 1% acetic acid solution (Sigma, 695092) for 5 mins. Dehydrated and mounted the slides as previous mentioned. Examined the slides using Leica DMi8 microscope.

**Cardiac MRI scan and analysis**

Cardiac MRI was performed at 1, 4, and 12 weeks (for pigs with 12-week follow-up) post-surgery. Replicates number for week 1 (n=10 medium control, n=10 CVP transplanted), week 4 (n=10 medium control, n=10 CVP transplanted) and week 12 (n= 3 medium control, n=5 CVP transplanted) weeks post cell-transplantation (refer to table 1). The scans were conducted on a 3.0T whole body MRI machine (Siemens Skyra, Siemens Medical Systems, Erlangen, Germany) with a standard cardiac flex coil (Siemens Medical Systems, Erlangen, Germany) as described[7, 8].

MRI imaging was performed by staff who was blinded to the animal groups. Pigs were sedated with intramuscular injection of ketamine/xylazine mixture.

Anesthesia was maintained with 2-2.5 % isoflurane on a ventilator after intubation. Heart rate and oxygen saturation were monitored throughout the examination. LV function was investigated using a segmented breath-held steady state free precession cine MRI imaging sequence. Contiguous 10 to 12 short-axis 2D slices without gap covering the LV from base to apex were acquired.

Global function was computed from the short-axis cine images by semi-automated segmentation of the LV endocardial and epicardial borders (from base to apex) at both end-diastole and end-systole using CVi42 analysis software (Circle Cardiovascular Imaging Inc., Canada). Infarct size was calculated from the late gadolinium enhancement images with scar surface area expressed as a percentage of the total LV surface area. Briefly, within the defined endocardial and epicardial borders, the LV area with signal intensity > mean+2SD of non-infarcted septal wall intensity, was considered as scar. Wall thickness (mm) was also measured from the software and results represented with 100 chords with AHA segmentations in a polar map.

**Computerized Tomography (CT) scan**

Whole body CT scans were acquired using a dual source 256 slices Siemens Somatom Definition Flash-CT camera. Animals were anesthetized, intubated, maintained on isoflurane and kept warm throughout the imaging procedure. Ketamine (100mg/kg + xylazine 20mg/kg, i.m.) was used for induction and 2-3% isoflurane was used for maintenance. Physiological monitoring were done on heart rate, $sPO_2$, and $CO_2$ %. The animals were kept warm by a convection warming blower, body temperature monitoring was done using a rectal probe. There was IV access for 0.9% NaCl drip and injecting contrast.

CT scans were done with the animal placed in supine position. An intravenous catheter was inserted into the ear vein and connected to a contrast injector. The scan area included the pig's snout superiorly and the lower limb inferiorly. There were 4 scan phases, pre-contrast, arterial phase (cardiac scan only), portal venous and delayed phase. Whole body scans were done in inspiration breath hold while scans over the chest were additionally ECG-gated. The contrast agent (Omnipaque 350®) was given at 1-3 ml/kg together with a saline bolus flush at 20 ml or 0.5 ml/Kg of the pig's weight depending on which was lower. Beta blockers (Esmolol 0.4mg/kg) was on hand to be used if the pig's heart rate went above 75 bpm to ensure the highest resolution of the cardiac images. Images were reconstructed and viewed on a SyngoVia workstation and analyzed to identify tumor formation in any organs. If a tumor was found, a necropsy will be performed when animals are sacrificed. The tumor will be explanted, fixed and sectioned to determine whether it was a teratoma or other tumor original from human cells by performing PCR for human Y chromosome detection.

**Electrophysiology Analysis**

Electromechanical mapping using 3D-NOGA system simultaneously registers the electrical and mechanical activities of the left ventricle, enabling assessment of myocardial viability. The aim of NOGA was to identify and localize arrhythmogenic foci and direct therapeutic procedures. This 3D mapping was done at weeks 2 and 4 by EP mapper and technical support team from Johnson and Johnson.

The external reference patch was placed on the pig's back, behind the heart. A 7 French introducer sheath was inserted into the right carotid artery, followed by the administration of 0.9 % NaCl/saline containing 1,000 IU of heparin. The fully deflected

NOGA-Star (Biosense Webster) catheter was advanced to the ascending aorta. The NOGA-Star catheter was moved through the aortic valve and the catheter tip oriented toward the apex. An apical point was obtained, followed by outflow tract, lateral, and posterior points, to form a 3D silhouette of the heart. The catheter was then deflected toward the anterior wall of the ventricle, during which multiple points of the anterior area were acquired with homogeneous distribution. The process was repeated to obtain additional reference points from the lateral, septal, anterior, and inferior walls, thus obtaining a full 3D map of the left ventricle.

During the mapping procedure, the NOGA workstation analyzed the 3D location of the catheter tip on the endocardial surface. The catheter tip moved in a stepwise manner to gather simultaneous unipolar and bipolar electrical signals for each mapping point. The stability of the catheter-to-wall contact was evaluated at every site in real time. After completing the map, postprocessing analysis filtered the unstable points caused by rhythm disturbances or internal cavities in the left ventricle. For all pigs in sinus rhythm, sinus rhythm left ventricular activation time mapping and voltage mapping was performed. For pigs with sustained ventricular tachycardia (VT), the VT was similarly mapped. VT was characterized as focal if there was radial spread of electrical activation from a point source.

Image and video postprocessing was performed with assistance by the team from Johnson and Johnson in a blinded fashion. Unipolar voltage maps indicates the exact positions of infarction. Unipolar voltage lower than 5 mV are considered scar (red color), unipolar voltage 5mV and more are considered healthy and viable myocardium (purple color). Local linear shortening (LLS) map provides information about wall movement (red = low movement, purple = healthy movement).

**Implantable cardiac monitoring system**

During the surgical procedure, Reveal LINQ$^{TM}$ cardiac monitoring system (Medtronic, USA) were subcutaneously placed in the left paraspinal area inferior to the angle of the scapula in pigs after surgery. The minimum sensed R wave post fixation of the Reveal LINQ was at least 0.3mV. Programmed settings were: sensitivity- 0.035mV, Blank-after-sense 150 ms, sensing-threshold decay 150 ms, ectopy rejection off. Recordings were intermittently downloaded once every 2 days. Detection criteria used were: Tachycardia – cycle length ≤ 370 ms (162bpm), number of intervals to detect (NID) 48 beats; Bradycardia – cycle length ≥ 2000ms (30 bpm), NID 4 beats; Pause – cycle length ≥ 3000ms. Every download was classified into three categories: (1) ventricular arrhythmia (defined as runs of 4 or more QRS complexes of different morphology compared to baseline or temporally adjacent sinus QRS together with evidence of atrioventricular dissociation), (2) bradycardia (defined as evidence of high grade atrioventricular block, sinus bradycardia ≤30 beats per minute and/or 3 or more sinus pauses or arrest with or without junctional escape rhythm within one recording), or (3) normal (downloads not belonging to the other two categories). Due to frequent T wave oversensing and QRS undersensing, all recordings were individually and manually read by an experienced electrophysiologist (Dr Eric Lim, NHCS). Equivocal recordings were adjudicated by a second electrophysiologist (Dr Paul Lim, NHCS).

**Quantification and statistical analysis**

Comparisons between groups were performed using two-way ANOVA with Tukey post hoc analysis unless otherwise stated. Values were reported as Mean ± SEM. P value < 0.05 % was considered statistically significant. Data were analyzed GraphPad Prism

(Version 9). Replicate information is indicated in the figures, figure legends and method details.

4. Yap LY, Li J, Phang IY, Ong LT, Ow JZ, Goh JC, Nurcombe V, Hobley J, Choo AB, Oh SK, Cool SM and Birch WR. Defining a threshold surface density of vitronectin

for the stable expansion of human embryonic stem cells. *Tissue engineering Part C, Methods*. 2011;17:193-207.

5. Bergenstrahle J, Larsson L and Lundeberg J. Seamless integration of image and molecular analysis for spatial transcriptomics workflows. *BMC Genomics*. 2020;21:482.

6. Yu G, Wang LG, Han Y and He QY. clusterProfiler: an R package for comparing biological themes among gene clusters. *OMICS*. 2012;16:284-7.

7. Ye L, Chang YH, Xiong Q, Zhang P, Zhang L, Somasundaram P, Lepley M, Swingen C, Su L, Wendel JS, Guo J, Jang A, Rosenbush D, Greder L, Dutton JR, Zhang J, Kamp TJ, Kaufman DS, Ge Y and Zhang J. Cardiac repair in a porcine model of acute myocardial infarction with human induced pluripotent stem cell-derived cardiovascular cells. *Cell Stem Cell*. 2014;15:750-61.

8. Ye L, D'Agostino G, Loo SJ, Wang CX, Su LP, Tan SH, Tee GZ, Pua CJ, Pena EM, Cheng RB, Chen WC, Abdurrachim D, Lalic J, Tan RS, Lee TH, Zhang J and Cook SA. Early Regenerative Capacity in the Porcine Heart. *Circulation*. 2018;138:2798-2808.
